# Putative target antigens of the stereotyped intrathecal B cell response in multiple sclerosis

**DOI:** 10.64898/2026.01.03.697481

**Authors:** Johanna H. Wagnerberger, Mari B. Gornitzka, Felix F. Loeffler, Jan Terje Andersen, Victor Greiff, Frode Vartdal, Andreas Lossius

## Abstract

Intrathecal antibody production is a hallmark of multiple sclerosis (MS), yet the antigenic targets of these antibodies remain elusive. Intrathecal antibody-secreting cells in MS patients characteristically produce IgG1 antibodies carrying the G1m1 allotype and preferentially use IGHV4 heavy-chain genes paired with IGKV1 or IGKV3 light chains. This stereotyped pattern points to a common antigen-driven selection process. To test this hypothesis, we generated monoclonal antibodies from intrathecal B-lineage cells bearing this stereotyped B-cell receptor configuration and screened them against a comprehensive library of human and Epstein-Barr virus proteins. In parallel, we developed and validated an immunoprecipitation-tandem mass-spectrometry (IPMS/MS) workflow for fresh-frozen brain tissue that preserves conformational epitopes. Several proteins, like CLDN11 and PLD1, were enriched by antibodies from multiple patients. Other targets, like MED21 and GIPC2, were unique to individual patients. Our study delivers a scalable pipeline for antigen discovery in MS and nominates candidate antigens for future validation.

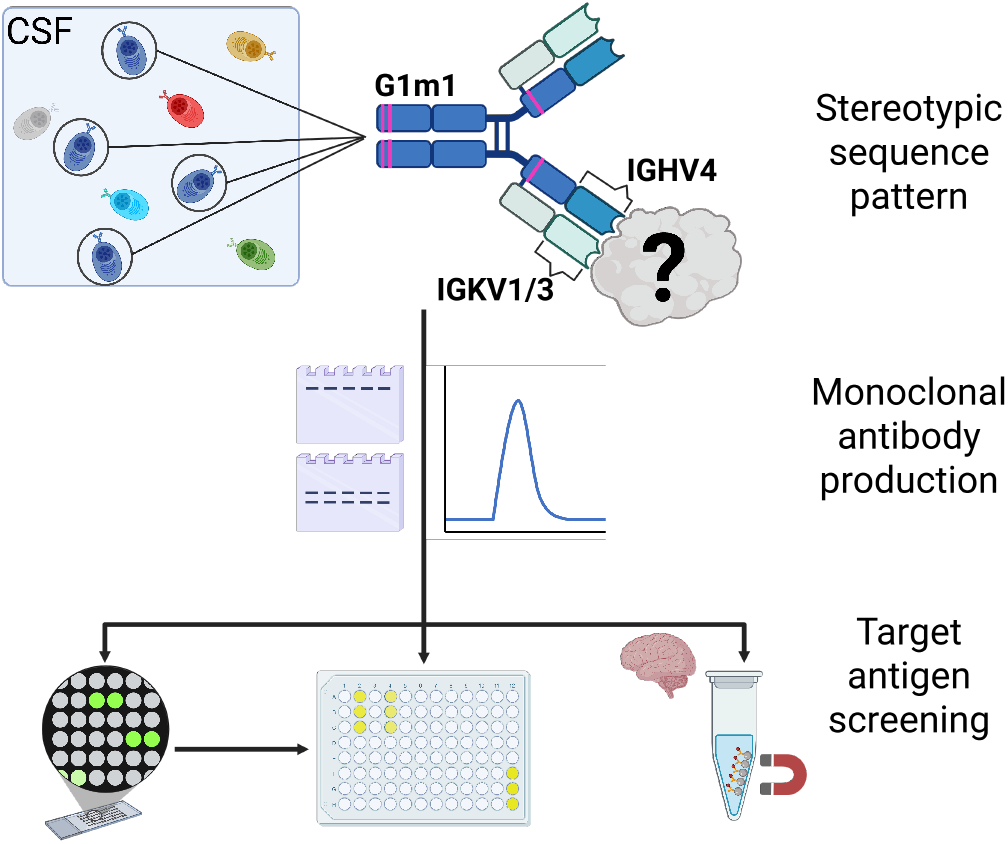

## Introduction

Multiple sclerosis (MS) is a chronic inflammatory demyelinating disorder of the central nervous system (CNS). A hallmark of MS is intrathecal synthesis of immunoglobulin G (IgG), first demonstrated by Kabat and colleagues in the late 1940s (Kabat, Glusman, and Knaub 1948). More sensitive techniques, such as isoelectric focusing, have later established that more than 90% of patients exhibit oligoclonal bands (OCBs) in their cerebrospinal fluid (CSF) (Dobson et al. 2013), indicating local IgG production by antibody-secreting cells (ASCs) (Obermeier et al. 2008; Polak et al. 2023). Notably, a similar CNS-restricted humoral immune response is also observed in chronic CNS infections (Owens et al. 2011) and in certain autoimmune encephalitides, including in a substantial proportion of patients with N-methyl-D-aspartate receptor (NMDAR) encephalitis (Blinder and Lewerenz 2019; Dalmau et al. 2008).

In chronic CNS infections, intrathecally produced IgG is directed against the causative pathogen. Antibodies isolated from the CSF of patients with subacute sclerosing panencephalitis, for example, primarily react to the disease-causing measles virus (Vandvik et al. 1976). Along the same line, ASCs targeting disease-specific autoantigens are enriched in the CSF of patients with NMDAR (Kreye et al. 2016), LGI1 (Kornau et al. 2020), and CASPR encephalitis (Theorell et al. 2024).

In contrast, targets of the intrathecal humoral immune response in MS remain elusive. Numerous candidate autoantigens have been proposed, yet none are widely accepted as disease-specific (Höftberger et al. 2022). One theory posits that the intrathecal antibody response represents a multi-target secondary reaction against cellular debris (Brändle et al. 2016; Grabar 1974). Another possibility is that these antibodies target an infectious agent. Given the strong epidemiological connection between Epstein-Barr Virus (EBV) infection and MS (Bjornevik et al. 2022; Thacker, Mirzaei, and Ascherio 2006), several studies have examined intrathecal production of EBV-reactive antibodies; however, results have been inconclusive (Owens et al. 2011; Sargsyan et al. 2010). Other groups have focused on antibodies that cross-react between EBV proteins and CNS autoantigens, hoping to uncover a molecular link between infection and autoimmunity (Tsai et al. 2017). Recently, Lanz et al. identified a monoclonal antibody that binds both EBV nuclear antigen 1 (EBNA1) and the human celladhesion molecule GlialCAM (Lanz et al. 2022), renewing interest in EBV-driven mechanisms in MS pathogenesis.

We previously applied full-length single-cell RNA sequencing to B-lineage cells isolated from the CSF of MS patients (Lindeman et al. 2022). The analysis revealed a conserved B-cell receptor (BCR) architecture across individuals, characterized by preferential VH:VL pairing associated with an allelic variation in the IgG1 heavy chain constant region (the G1m1 allotype). Notably, G1m1 BCRs frequently used IGHV4-family genes and showed the highest rate of pairing with IGKV1 and IGKV3 light-chain families. Such restricted variable-gene usage is a hallmark of antigen-driven selection and has been reported in other settings, including celiac disease (Di Niro et al. 2012), HIV infection (Gorny et al. 2009), influenza (Ekiert et al. 2009), and COVID-19 (Kim et al. 2021), where BCR repertoires are shaped by specific antigenic epitopes.

In this study, we aimed to identify a common antigenic target recognized by the stereotyped B-cell response in MS. To this end, we selected CSF B-lineage cells with the stereotyped BCR motifs and recombinantly expressed their receptors as monoclonal antibodies (mAbs). These mAbs were screened against human and EBV proteins. We also developed and validated an immunoprecipitation protocol coupled with tandem mass spectrometry (IP-MS/MS) compatible with fresh-frozen human brain tissue, enabling sensitive detection of membrane proteins.

## Materials and methods

### Antibody selection

Antibody sequences were selected from single-cell sequencing data that contained the paired, full-length BCR sequences of CSF-residing B cells from 21 MS patients (Lindeman et al. 2022). The study was approved by the Committee for Research Ethics at the South-Eastern Norwegian Heath Authority (2009/23), and all patients had signed a written informed consent. The BCR sequences were filtered according to heavy-chain allele usage, light-chain pairing, V-gene usage, and the degree of clonal expansion. For patients where mass spectrometry data was available (MS2, MS8, MS10, MS14, and MS19) this information was utilized to select BCR sequences that matched to intrathecally synthesized IgG. All selected BCR sequences were part of an intrathecally expanded clone, except MS14_165, MS20_14, and MS21_115. The resulting 24 antibodies are all derived from BCR sequences that were G1m1, carried variable heavy chain genes of the IGHV4 family, and paired with kappa light chains carrying either IGKV1 or IGKV3.

From the literature, sequences of the antibodies “MS39p2w174” (Lanz et al. 2022) and “#008-218” (Kreye et al. 2016) with characterized target epitopes were selected as positive controls. As a negative control, we selected an antibody sequence from a naïve B cell found in the CSF of a G1m3 homozygous MS patient of our cohort (Lindeman et al. 2022).

### Recombinant expression of monoclonal antibodies

Selected antibody heavy and light chain variable sequences were synthesized by GenScript (Rijswijk, Netherlands). The heavy chain sequences were cloned into a modified pFUSE-CHIg plasmid, which carried one of two allotypes in the heavy chain. G1m17 and G1m1 was used for MS39p2w174 and the MS-derived mAbs with stereotyped patterns, whereas G1m3 and nG1m1 was used for #008-218 and the negative control antibody MS5_NB2. The variable segments of the light chains were cloned into pFUSE2-CLIg. Subsequently, Expi293F cells were transfected using the Gibco Expi293 Expression System (Thermo Fisher Scientific, Waltham, MA, USA) according to the manufacturer’s instructions (heavy-to light chain plasmid ratio 1:2). The cell media was harvested 5-6 days after transfection, and antibodies were purified using a HiTrap Protein A HP antibody purification column (Cytiva, Uppsala, Sweden). Purified antibodies were concentrated and buffer-exchanged to PBS containing 0.02% sodium azide using centrifugal filter columns with a 50 kDa cutoff (Merck Millipore, Burlington, MA, USA). Aliquots were subsequently stored at 4 °C.

### Antibody Quality control

Quality control of recombinantly expressed antibodies included assessment of purity and monomeric form by reducing and non-reducing SDS-PAGE and analytical size-exclusion chromatography (SEC). For SDS-PAGE, samples were mixed 1:5 with 6× non-reducing Laemmli sample buffer (Thermo Fisher Scientific, Waltham, MA, USA); *β*-mercaptoethanol was added to 1% (v/v) for reducing samples. After heating at 96°C for 10 min (non-reducing) or 15 min (reducing), samples were loaded onto precast Criterion TGX gels, 10% (non-reducing) or 12% (reducing) (Bio-Rad laboratories, Hercules, CA, USA). SEC was performed on purified mAbs (>2 mg/mL); 15 µL was injected on a Superdex 200 Increase 3.2/300 small-scale SEC column (Cytiva, Uppsala, Sweden), at 0.05 mL/min. In selected mAbs, plasmid DNA and antibody amino acid sequences were verified by sanger sequencing and LC-MS/MS, respectively. For LC-MS/MS, 20 µg of purified mAb was reduced, alkylated, digested by trypsin, desalted, and analyzed on an EVOSEP liquid chromatography system coupled to a timsTOFfleX or timsTOF-pro2 mass spectrometer (Bruker Daltonics, Bremen, Germany).

### Functional verification of control antibodies

For control antibodies, biding to the cognate target was confirmed. Binding of #008-218 to GluN1 was verified by immunocytochemistry on GluN1-transfected cells using a 4-point dilution series of #008-218 (1 - 500 µg/mL) and comparison to verified positive and negative control sera. Targets of MS39p2w174 were verified by ELISA on clear flat-bottom 96-well plates (Thermo Fisher Scientific, Waltham, MA, USA). We coated the plates with either EBNA-1_AA386-405_ or glialCAM_AA370-389_ (both peptides from GenScript, Rijswijk, Netherlands) in two-fold dilution series. After each step, four washes with PBST were performed. MS39p2w174 was applied at 2 µg/mL, followed by a goat anti-human IgG-alkaline phosphatase secondary antibody (A18832, Thermo Fisher Scientific, Waltham, MA, USA) and AP substrate (S0942, Sigma Aldrich, MO, USA). Absorbance at 405 nm was recorded on a FLUOstar Omega microplate reader (BMG Labtech, Ortenberg, Germany).

### Proteome array

The antibodies were screened in pools of five antibodies per HuProt Human Proteome Microarray v4.0 (CDI Labs, Puerto Rico). The pools were sent to PEPperPRINT GmbH (Heidelberg, Germany), where the HuProt Chip screening was performed. Briefly, background interactions to the proteome array were first determined by incubating the chip with secondary goat anti-human IgG (H+L) DyLight680 antibody and measuring the fluorescent readout on an Innopsys InnoScan 710-IR Microarray Scanner. Subsequent incubation with the MS antibody pools was performed at a concentration of 50 µg/mL (10 µg/mL per antibody) before washing and repeating the staining with the same secondary antibody. Grid alignment and quantification of spot intensities were performed using Mapix 9.1.0 (Innopsys), and R 4.1.2 was used to calculate background-corrected spot intensities. Common hits among more than one pool were filtered for proteins known to interact with other parts of the antibody than the variable region before further investigating the hits.

### ELISA

From the HuProt assay, protein spots with high background-corrected fluorescence were further investigated by ELISA in-house. The proteins in question (Supplementary table S1) were diluted to 2 µg/mL in PBS and incubated on a 96-well polystyrene plate overnight. The following day, the plate was washed with PBST (0.05% Tween20) and blocked with PBST-B (1% BSA) before each MS-derived antibody (2 µg/mL) was incubated on the plate in triplicate. Antibody binding was verified using alkaline phosphatase-conjugated goat anti-human Fc as a secondary antibody (A18832, Invitrogen, MA, USA). After washing the plate, Phosphatasesubstrate (S0942, Sigma Aldrich, MO, USA) diluted to 1 mg/mL in diethanolamine buffer was added, and absorbance at 405nm was measured 15min later using a FLUOstar Omega microplate reader (BMG Labtech, Ortenberg, Germany).

### EBV microarray

The PEPperCHIP®Epstein-Barr Virus Peptide Microarray (PEPperPRINT GmbH) was acclimated to room temperature before assembling the incubation tray. The MS-derived antibodies were screened against the microarray in pools of five and incubated overnight as described by the manufacturer. The control antibody MS39p2w174 was incubated in a second round after the MS-derived antibodies had already been screened on the same chip. MS39p2w174 was incubated for two hours at 4 °C, 160rpm shaking. Antibody concentration during all incubations was 10 µg/mL. Binding was measured by fluorescent intensity using anti-human Fc conjugated to DyLight 650 (Thermo Fisher Scientific, Waltham, MA, USA).

### Immunoprecipitation

Six brain tissue blocks containing plaques from three individuals diagnosed with MS were provided by the Netherlands Brain Bank (Amsterdam, Netherlands). For each round of experiments, approximately 100mg of tissue was cut from each tissue block and placed in 500 µL lysis buffer (25 mM HEPES, 150 mM NaCl, 2 mM EDTA, 1.5% (w/v) n-Dodecyl-*β*-Maltoside (DDM) and 1× Halt Protease and Phosphatase Inhibitor Cocktail). The buffer volume was adjusted to 10 µL per mg tissue after mincing the tissue using five strokes of a micropestle. Subsequently, each sample was sonicated for 7×10 sec (100 watts, 100% amplitude), with a 30 sec cooling period on ice between each sonication round. The resulting lysate was incubated on rotation (4 °C, 20 min) before insoluble particles were pelleted by centrifugation (4 °C,16.000 rcf, 20 min). The supernatant was kept cool while the pellet was reprocessed in 500 µL lysis buffer by sonicating the pellet for 2×10 sec. After a second centrifugation of the supernatants and reprocessed pellets (4 °C,16.000 rcf, 30 min), all the supernatants were pooled into one lysate master mix.

The protein concentration of the brain lysate was measured on a 1:10 dilution using the Pierce BCA Protein Assay kit according to the manufacturer’s instructions. If the protein concentration was above 3.0 mg/mL, the lysate was diluted to this concentration before pre-clearing the lysate from endogenous immunoglobulins by incubating it with Dynabeads protein A (30 min, 4 °C). Subsequently, the lysate was divided into 520 µL aliquots.

The final immunoprecipitation was performed by adding 20 µg of mAb to a lysate aliquot and incubating it on rotation overnight at 4 °C. The following day, 100 µL Dynabeads protein A were added and incubated for 30 min at room temperature. The Dynabeads were subsequently separated from the lysate using a magnet and washed twice with strong wash buffer (25 mM HEPES, 150 mM NaCl, 2 mM EDTA, 0.5% (w/v) DDM) and three times with mild wash buffer (0.05% (w/v) DDM). The antibody-antigen complexes were then reduced, alkylated, and digested directly on the bead by transferring the beads to 100 µL 50 mM NH4HCO3 and adding 1 µL 1 M Dithiothreitol (incubated at 56 °C for 30 min), 2.7 µL 550 mM Iodoacetamide (incubated protected from light for 45 min) and 1 µL of 1 µg/µL trypsin (incubated at 37 °C overnight). Peptide desalting was performed using C18 stage tips. This procedure was performed in triplicate for all monoclonal antibodies.

Subsequent mass spectrometry was performed on an EVOSEP liquid chromatography system coupled to a timsTOFfleX or timsTOF-pro2 mass spectrometer (Bruker Daltonics, Bremen, Germany) via a CaptiveSpray ion-source. Mass spectra for MS and MS/MS scans were recorded between m/z 100 and 1700. Data-dependent acquisition was performed using 10 PASEF MS/MS scans per cycle with a near 100% duty cycle.

Protein identification was performed using MaxQuant v2.6.0.0 (Cox and Mann 2008) and a human proteome reference downloaded from UniProt (April 2024). The resulting protein groups file was then used in Perseus v2.0.11 (Tyanova et al. 2016) to filter the results and compare protein LFQ values across samples. Further filtering of the protein groups file was done by removing potential contaminants, reverse hits, and proteins only identified by peptides carrying modified amino acids (“only identified by site”). Next, the proteins enriched by MS-derived antibodies were compared to the negative control antibody MS5_NB2. The LFQ values were log2-transformed, and only proteins that contained valid LFQ values in at least two triplicates were kept for further analysis. After replacing missing values, a two-sample t-test (threshold p-value 0.05) was used to compare the LFQ values of the respective mAb to MS5_NB2. To identify and filter out non-specific binding, we performed the immunoprecipitation protocol using an anti-measles nucleocapsid mAb “2B4” (Burgoon et al. 1999; Owens Gregory P. et al. 2006) and an anti-SARS-CoV-2 spike protein mAb “P05DHu” (Thermo Fisher Scientific, Waltham, MA, USA) and excluded proteins enriched by either of these antibodies from further analysis. The remaining significantly enriched proteins were visualized as volcano plots using the tidyverse package (v2.0.0) in R v4.3.2. An overall summary of proteins enriched by multiple MS mAbs was produced in R by using the UpSetR package (v1.4.0) in addition to tidyverse.

## Results

### Selection and quality control of monoclonal antibodies

From 2,165 B-lineage cells profiled by full-length single-cell RNA-sequencing across 21 MS patients (Lindeman et al. 2022), 375 ASCs met our predefined stereotyped pattern criteria (see Methods). These 375 ASCs were derived from 13 MS patients (Table 1). We selected twenty-four representative BCR sequences from this pool for recombinant monoclonal antibody (mAb) production (Table 2). Because most of the chosen BCRs derived from expanded clonal families, these 24 unique mAbs correspond to 73 single ASCs whose BCRs are identical or clonally related to one of the mAbs.

**Table 1.**
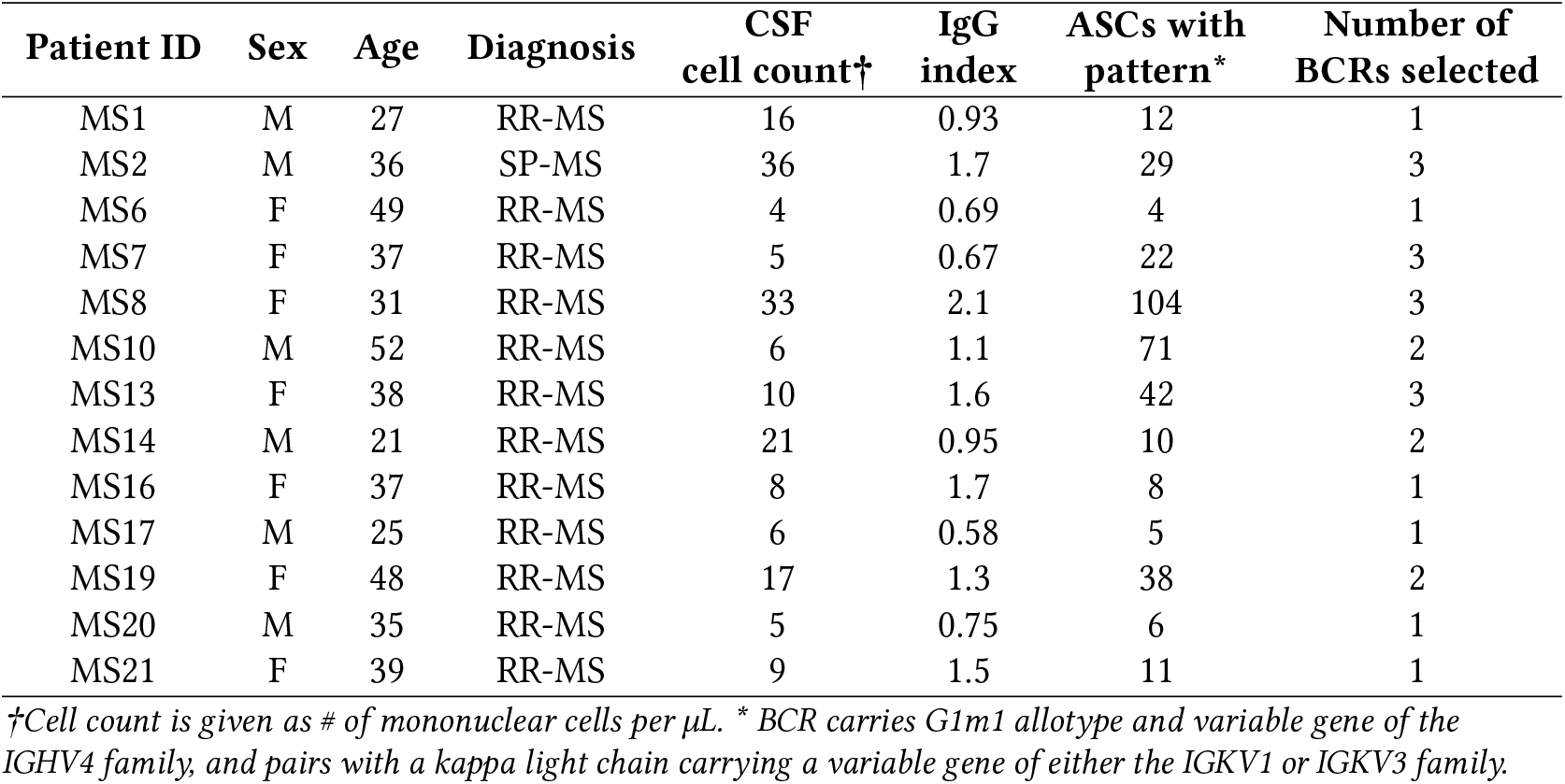
Patient information.

**Table 2.**
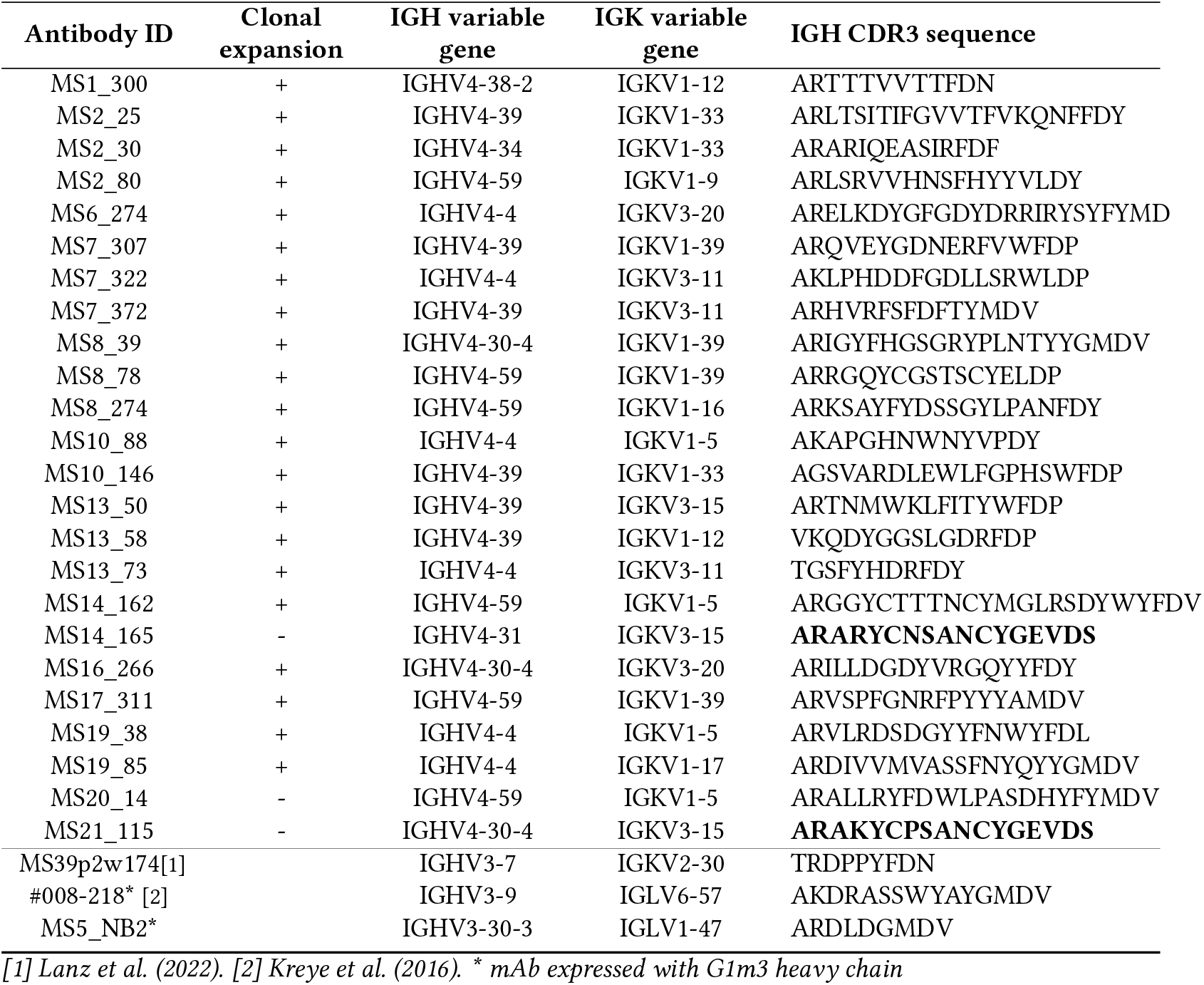
Monoclonal antibody sequence information, including whether the parent ASC was part of an expanded clone. CDR3 sequences that were matched in a similarity search are indicated in bold. The bottom three rows display control antibody information, with two of them not carrying the G1m1 allotype (indicated by a star).

Quality control of the recombinantly expressed mAbs consisted of methods verifying antibody purity (Figure 1A), confirming sequence identity (Figure 1B), and ensuring monomeric form (Figure 1C; a summary table of all mAbs can be found in Supplementary Table S2). Two well-characterized monoclonal antibodies were included as controls and produced in-house in the same expression system as the MS mAbs. Antibody #008-218 binds a conformational epitope on the GluN1 subunit (gene name GRIN1) of the NMDA receptor (Figure 1D) and was originally isolated from the CSF of an anti-NMDAR encephalitis patient (Kreye et al. 2016). Antibody MS39p2w174 was identified in the CSF of an MS patient by Lanz et al. (2022). It targets a linear epitope with sequence similarity between EBNA1 and GlialCAM (gene name HEPACAM; Figure 1E). After undergoing the same quality control pipeline used for the MS-derived mAbs, both control antibodies were functionally validated. Antibody #008-218 bound its target in a cell-based assay with GluN1-overexpressing cells (Figure 1F), whereas antibody MS39p2w174 recognized synthetic peptides from both EBNA1 and GlialCAM in ELISA (Figure 1G).

**Figure 1.**
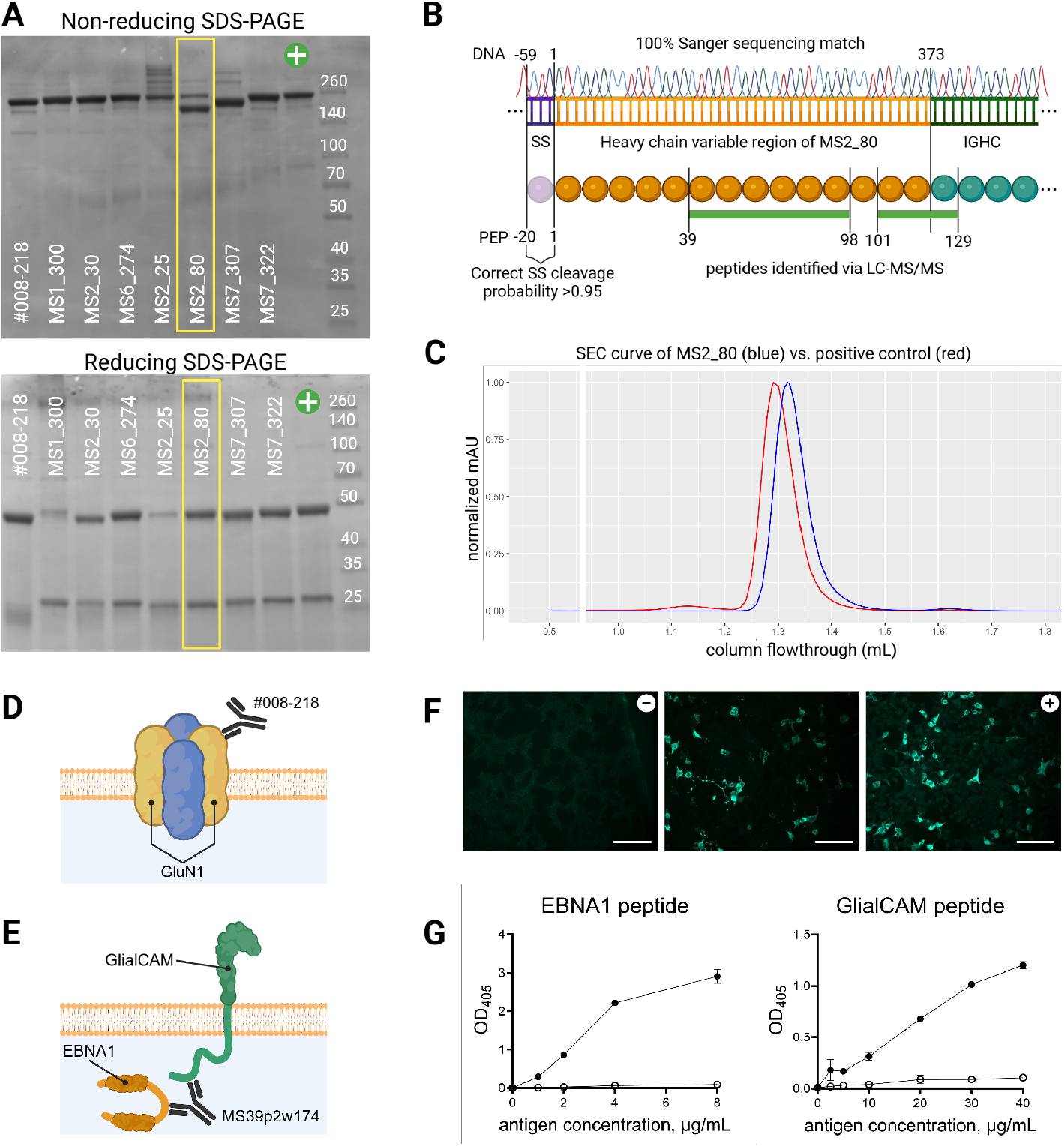
Quality control of in-house antibody production. (A) Antibodies were screened for size and purity on SDS-PAGE; a purified myeloma IgG served as a positive control (+). (B-C) Example of further quality control for selected antibodies, illustrated with mAb MS2_80: (B) plasmid Sanger sequencing, protein mass-spectrometry, and in silico signal sequence (SS) cleavage prediction; (C) size-exclusion chromatography (SEC). (D-E) Schematic of control antibodies and their targets: (D) #008-218 (anti-GluN1) and (E) MS39p2w174 (anti-EBNA1/GlialCAM). (F) Binding of #008-218 to cells overexpressing GluN1 was verified alongside negative (-) and positive (+) control sera. (G) Titration of MS39p2w174 (filled circles) and an isotype control (hollow circles) against increasing concentrations of EBNA1_AA386-405_ and GlialCAM_AA370-389_, assessed by ELISA.

### Array-based screening methods reveal no common targets

Since EBV is closely linked to MS epidemiologically, we screened all MS mAbs against a peptide microarray containing approximately 5 500 overlapping 13-mer peptides spanning over 18 EBV proteins. This confirmed that MS39p2w174 bound to its published linear EBNA1-epitope; however, none of our MS mAbs showed marked reactivity towards any of the screened EBV peptides (Supplementary Figure 1).

To investigate human targets, the MS mAbs were pooled in groups of five and screened against the HuProt Human Proteome Microarray, which covers over 20 000 full-length proteins. This comprehensive screen revealed two strong signals: one against GIPC2 in Pool 1 and the other against MED21 in Pool 2 (Figure 2A). Subsequent ELISAs with the individual mAbs from these pools confirmed that MS19_38 binds GIPC2 and MS13_50 binds MED21 (Figure 2B). A few protein targets were detected across multiple pools (Figure 2A); however, their microarray signals were only low to intermediate and failed orthogonal validation by ELISA (data not shown). Notably, although HuProt internal controls showed robust signals, our in-house positive control (#008-218; anti-GluN1) did not bind its target on the array (Figure 2A). This demonstrates that disease-specific key epitopes may have conformational requirements that are not available for screening on the microarray.

**Figure 2.**
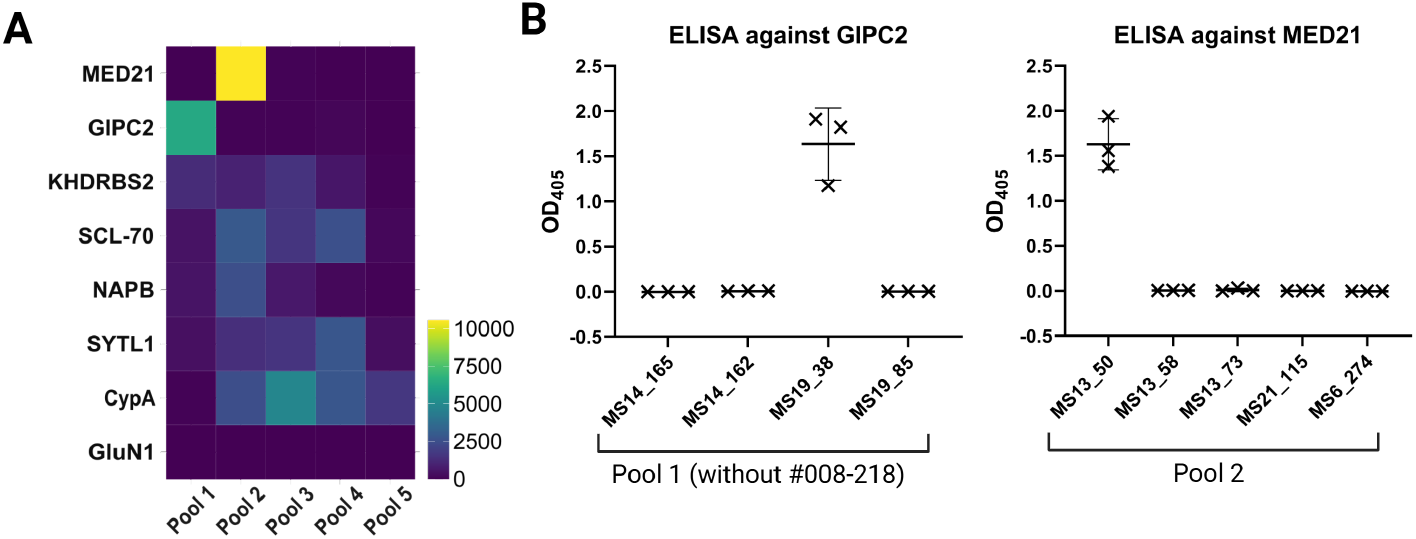
HuProt screening and subsequent investigation of hits. (A) Background-corrected fluorescent intensities on the HuProt microarray for pooled MS-derived mAbs (five mAbs per pool). With the exception of the known target of #008-218 (GluN1), the displayed proteins showed either high binding to a single pool or moderate binding across multiple pools. No protein displayed high binding across multiple pools. (B) Background-corrected OD_405_ of subsequent ELISA experiments confirmed GIPC2 and MED21 as targets of a single antibody in Pool 1 and Pool 2, respectively.

### An enhanced protocol of immunoprecipitation conserves native protein structures

The observation that #008-218 did not identify its antigen on a protein microarray, even though it bound well to cells transfected with its target (Figure 1E), led us to consider methods that better preserve native protein conformations during the screen. This naturally led to the use of fresh-frozen brain tissue, as proteins in the diseased organ would be in correctly folded/naturally relevant conformations.

We focused on producing a protocol to preserve and solubilize membrane-bound proteins for IP. We chose a HEPES-based lysis buffer with the detergent n-Dodecyl-*β*-Maltoside as a solubilizing agent and no added denaturing agents. Using this protocol on mouse cerebellum, we successfully enriched proteins of the NMDA receptor complex using the mAb #008-218 and GlialCAM using mAb MS39p2w174 (Figure 3A). This was also replicated in human brain lysate (Supplementary Figure 2).

**Figure 3.**
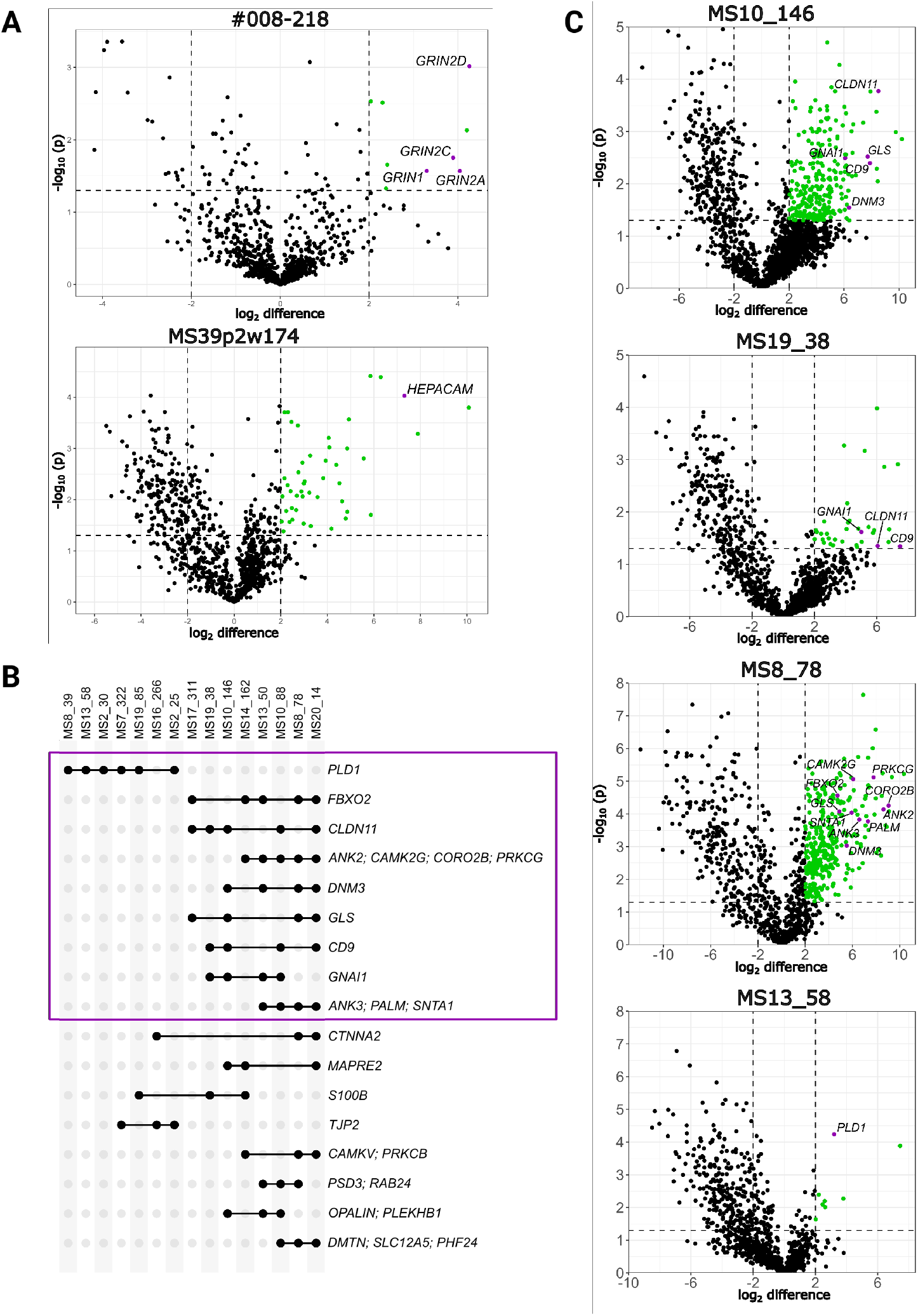
Immunoprecipitation of multiple sclerosis brain tissue. (A) Volcano plots from IP-MS/MS on mouse cerebellum show that control antibodies enrich their cognate antigen along with known interacting proteins (purple). (B) Curated list of proteins enriched by three or more MS mAbs in IP-MS/MS of MS patient white matter. (C) Representative volcano plots displaying the proteins in (B) that are enriched by four or more antibodies (highlighted in purple).

### Shared and private candidate targets identified by immunoprecipitation of MS brain tissue

Each MS mAb was used individually for IP of lysates from MS brain tissue (see Methods). Plaque-containing tissue was lysed, and IP was directly followed by tryptic digestion for LC-MS/MS. Raw data were processed with MaxQuant and known contaminants/background were filtered as described in Methods. Because these mAbs were selected for a shared stereotyped pattern–suggestive of a common antigen–we focused on proteins significantly enriched by at least three antibodies from more than one patient, yielding 84 proteins (Supplementary Table S3). To evaluate potential nonspecific interactions, we queried the CRAPome database (Mellacheruvu et al. 2013) to determine how frequently each candidate appears as a contaminant in affinity purification-mass spectrometry datasets. We then examined brain protein expression and celltype stratified RNA profiles for the individual proteins using the Human Protein Atlas (Thul et al. 2017) (Supplementary Table S4). Figure 3B shows the final set of 17 candidate proteins of interest, and representative volcano plots for targets shared by four or more antibodies are shown in Figure 3C.

In addition to the shared antigen candidates, some mAbs precipitated distinct proteins or protein complexes. Four prominent examples of these “private” targets are shown in Supplementary Figure 3. Notably, the two individual targets identified using the proteome microarray and ELISA (Figure 2B) were not confirmed by immunoprecipitation.

## Discussion

In this study, we used a stereotyped amino acid motif to select intrathecal BCRs from MS patients and recombinantly expressed them as mAbs. We screened these mAbs against thousands of human and EBV proteins using protein microarrays. In parallel, we developed an IP-MS/MS protocol compatible with fresh-frozen human brain tissue and sufficiently sensitive to preserve and enrich membrane proteins. As proof of concept, the protocol successfully precipitated the GluN1 subunit of the NMDAR when probed with a conformationally dependent anti-GluN1 antibody (Kreye et al. 2016) and confirmed the GlialCAM-reactive MS mAb reported by Lanz et al. (2022).

Since our approach selected mAbs that shared the same stereotyped pattern, we primarily sought to identify a target antigen common across patients. Using the IP-MS/MS protocol, we detected several proteins precipitated by at least three mAbs from more than one individual. Among these was claudin-11 (CLDN11), a multi-pass tight junction protein enriched in CNS myelin that has been proposed as an autoantigen in MS (Höftberger et al. 2022; J. M. Bronstein et al. 1999; Quintana et al. 2008). CLDN11 can account for up to 7% of total myelin protein (J. Bronstein, Micevych, and Chen 1997) and was immunoprecipitated by five mAbs. In four of the five cases, CLDN11 coimmunoprecipitated with the tetraspanin CD9, consistent with known claudin-tetraspanin interactions (Kovalenko, Yang, and Hemler 2007). An early study reported CLDN11-reactive CSF antibodies in approximately 80% of MS patients (J. M. Bronstein et al. 1999), whereas a later analysis using native protein found no MS-specific response (Aslam et al. 2010). These discrepancies underscore the need for independent validation of the shared-antigen list generated here.

The HuProt screen did not reveal any antigens shared by multiple mAbs, but it did produce strong and specific signals for MED21 and GIPC2, each recognized by a single mAb from different patients. MED21 is a subunit of the mediator complex that regulates RNA polymerase II activity (Sato et al. 2016), whereas GIPC2 is believed to act as an intracellular adaptor molecule bridging myosin VI with transmembrane proteins (Katoh 2013). Both targets were independently confirmed by ELISA, yet were not reproduced by IP-MS/MS in brain tissue. For GIPC2, the simplest explanation is its low abundance in adult white matter (Thul et al. 2017). For MED21, epitope masking within the intact mediator complex might be more likely: the protein is part of the central module and adopts several conformations depending on its binding partners (Tsai et al. 2017). It is thus highly likely that the conformation and accessible sites of the isolated MED21 protein on HuProt/ELISA and the naturally occurring complex in the lysate differ. The importance of native context is underscored by our control mAb #008-218, which readily precipitates GluN1 from brain lysate yet fails to bind the subunit on the microarray. Notably, comparable losses of reactivity in the HuProt assay have been reported for serum and CSF antibodies in anti-NMDAR encephalitis patients and other patients with known neurological antigens (McKeon et al. 2023). Taken together, our observations suggest that the single-antibody HuProt hits in our screening most likely recognize epitopes exposed only in isolated or partially unfolded proteins, whereas antigens that rely on native quaternary structure or a membrane context are preferentially recovered by IP-MS/MS.

Like MED21 and GIPC2, many of the proteins enriched by multiple mAbs in our IP-MS/MS experiments are intracellular. One example is phospholipase D1 (PLD1), an enzyme implicated in signal transduction and neurite outgrowth (Klein 2005). This bias toward intracellular targets could simply reflect their greater abundance in the lysate, but we cannot exclude the possibility that part of the humoral response in MS may be directed against antigens normally sequestered inside cells. Brändle et al. (2016) reported a similar pattern when analyzing mAbs derived from isolated OCBs and proposed that such reactivity could arise secondary to tissue destruction, which releases otherwise hidden intracellular proteins. Several previously proposed MS autoantigens, like *αβ*-crystallin (Rand et al. 1998; Thomas et al. 2023) and neurofilament light chain (Puentes et al. 2017), are likewise intracellular, although they were not detected in our study.

A linear EBV peptide screen revealed no binding to our panel of mAbs. It did, however, confirm the published EBNA1 epitope of MS39p2w174 (Lanz et al. 2022). None of our MS mAbs showed any reactivity towards this epitope. It is often reported that MS patients have stronger antibody reactivity towards specific EBNA1 peptides (Hecker et al. 2016; Ruprecht et al. 2014; Sattarnezhad et al. 2025), and the epitope of MS39p2w174 is within one such peptide. However, most of these studies have not corrected for higher antibody reactivity towards the entire EBNA1 protein. A recent report by Cortese et al. (2024) reports no significant reactivity towards any singular peptide after such a correction, weakening the hypothesis that one specific cross-reactivity drives the disease. Since EBV is a B-cell tropic virus (Sathiyamoorthy et al. 2016), it is also possible that the connection between viral infection and disease does not lie in the antigenic target of the BCR, but rather in how the viral infection alters its host. Most studies, including this one, have primarily assessed reactivity towards linear EBV epitopes. However, it could be that there are unexplored reactivities toward conformational EBV targets that hold the key to MS development and/or progression.

Identifying conformational epitopes, especially on multi-pass membrane proteins, is notoriously difficult in high-throughput screens, because the physiochemical conditions required to preserve native folding differ from protein to protein and often conflict (Choy et al. 2021). The IP-MS/MS workflow developed here, on the other hand, appears to retain both membrane topology and higher-order complexes. This is best exemplified by its ability to precipitate several subunits of the NMDA receptor from mouse cerebellum. The GluN1 subunit, containing the target epitope, was again enriched in human brain lysate (Supplementary Figure 2), although the auxiliary GluN2/3 subunits did not reach significance. One of the MS mAbs pulled down the full pyruvate dehydrogenase (PDH) complex (Supplementary Figure 3), showing that large mitochondrial protein complexes can also be preserved. Likewise, the cytosolic enzyme *δ*-aminolevulinate dehydratase (ALAD), enriched by another MS mAb, displayed a very high log2-enrichment score in its sample, consistent with the retention of its native homo-octameric/hexameric structure (Breinig et al. 2003).

It should be noted that a prerequisite for success in the immunoprecipitation experiment is sufficient availability of the putative target in the MS lesion or surrounding normal-appearing white matter. This experiment, thus, cannot be interpreted as a screen of the human proteome. Additionally, low-affinity interactions might not be identifiable in this approach, as immunoprecipitation generally requires interactions of moderate to high affinity if no additional crosslinking methods are employed (Colwill and Gräslund 2011; Dyson et al. 2011). Some of our MS mAbs enriched very few proteins past the significance threshold, despite the high sensitivity of LC-MS/MS. This could indicate that the target was not present in the lysate or that the affinity between antibody and antigen was not high enough to withstand subsequent washing and shearing forces. The mass-spectrometry analysis would also not be able to detect non-protein targets like phospholipids and gangliosides, both of which have been proposed as antibody targets in MS patients previously (Höftberger et al. 2022).

In conclusion, we present a scalable workflow for antigen discovery in MS, which is also widely applicable for other CNS diseases and infections. Our immunoprecipitation protocol preserves conformational epitopes and intact protein complexes, as shown by the pull-down of the NMDAR complex with a conformation-dependent anti-GluN1 mAb. Applying the method to stereotyped MS mAbs uncovered several candidate antigens shared across patients, which now warrant targeted validation in future studies. However, no single protein stood out as commonly enriched by all mAbs, and several mAbs precipitated distinct targets, pointing to greater diversity than initially hypothesized. This highlights the need for further studies to clarify not only the target of intrathecal antibodies in MS, but also how and why they appear in the CNS.

## Acknowledgements

We would like to thank Dr. Jakob Kreye and Dr. Harald Prüß for providing the sequence of their anti-NMDAR antibody “#008-218”, as well as Dr. Gregory Owens for providing the sequence of the anti-measles antibody “2B4”. They were invaluable controls in our experiments. We also thank Anne Mattison Lien at the section of medical immunology, Oslo university hospital Ullevål for screening #008-218 against GluN1-transfected cells. The Proteomics core facility at Oslo University Hospital, particularly Dr. Maria Ekman Stensland, provided essential insight for protocol development and result analysis of the IP-MS/MS data. We are deeply grateful to the individuals with MS who donated cerebrospinal fluid for monoclonal antibody generation, and to the three donors and their families for post-mortem brain tissue provided through the Netherlands Brain Bank. Their generosity made this work possible.

## References

Aslam, Muhammad et al. (Apr. 2010). “The antibody response to oligodendrocyte specific protein in multiple sclerosis”. eng. In: Journal of Neuroimmunology 221.1-2, pp. 81–86.

Bjornevik, Kjetil et al. (Jan. 2022). “Longitudinal analysis reveals high prevalence of Epstein-Barr virus associated with multiple sclerosis”. In: Science 375.6578. Publisher: American Association for the Advancement of Science, pp. 296–301.

Blinder, Tetyana and Jan Lewerenz (July 2019). “Cerebrospinal Fluid Findings in Patients With Autoimmune Encephalitis—A Systematic Analysis”. In: Frontiers in Neurology 10, p. 804.

Brändle, Simone M. et al. (July 2016). “Distinct oligoclonal band antibodies in multiple sclerosis recognize ubiquitous self-proteins”. eng. In: Proceedings of the National Academy of Sciences of the United States of America 113.28, pp. 7864–7869.

Breinig, Sabine et al. (Sept. 2003). “Control of tetrapyrrole biosynthesis by alternate quaternary forms of porphobilinogen synthase”. en. In: Nature Structural & Molecular Biology 10.9. Publisher: Nature Publishing Group, pp. 757–763.

Bronstein, J. M. et al. (July 1999). “A humoral response to oligodendrocyte-specific protein in MS: a potential molecular mimic”. eng. In: Neurology 53.1, pp. 154–161.

Bronstein, J.m., P.e. Micevych, and K. Chen (1997). “Oligodendrocyte-specific protein (OSP) is a major component of CNS myelin”. en. In: Journal of Neuroscience Research 50.5, pp. 713–720.

Burgoon, M.P et al. (Feb. 1, 1999). “Cloning the antibody response in humans with inflammatory CNS disease: isolation of measles virus-specific antibodies from phage display libraries of a subacute sclerosing panencephalitis brain”. In: Journal of Neuroimmunology 94.1, pp. 204–211.

Choy, Brendon C. et al. (Mar. 2021). “A 10-year metaanalysis of membrane protein structural biology: Detergents, membrane mimetics, and structure determination techniques”. In: Biochimica et Biophysica Acta (BBA) - Biomembranes 1863.3, p. 183533.

Colwill, Karen and Susanne Gräslund (July 2011). “A roadmap to generate renewable protein binders to the human proteome”. en. In: Nature Methods 8.7. Publisher: Nature Publishing Group, pp. 551– 558.

Cortese, Marianna et al. (May 2024). “Serologic Response to the Epstein-Barr Virus Peptidome and the Risk for Multiple Sclerosis”. In: JAMA Neurology 81.5, pp. 515–524.

Cox, Jürgen and Matthias Mann (Dec. 2008). “MaxQuant enables high peptide identification rates, individualized p.p.b.-range mass accuracies and proteome-wide protein quantification”. en. In: Nature Biotechnology 26.12. Publisher: Nature Publishing Group, pp. 1367–1372.

Dalmau, Josep et al. (Dec. 2008). “Anti-NMDA-receptor encephalitis: case series and analysis of the effects of antibodies”. In: The Lancet Neurology 7.12, pp. 1091–1098.

Di Niro, Roberto et al. (Feb. 2012). “High abundance of plasma cells secreting transglutaminase 2-specific IgA autoantibodies with limited somatic hyper-mutation in celiac disease intestinal lesions”. eng. In: Nature Medicine 18.3, pp. 441–445.

Dobson, Ruth et al. (Aug. 2013). “Cerebrospinal fluid oligoclonal bands in multiple sclerosis and clinically isolated syndromes: a meta-analysis of prevalence, prognosis and effect of latitude”. en. In: Journal of Neurology, Neurosurgery & Psychiatry 84.8. Publisher: BMJ Publishing Group Ltd Section: Multiple sclerosis, pp. 909–914.

Dyson, Michael R. et al. (Oct. 2011). “Mapping protein interactions by combining antibody affinity maturation and mass spectrometry”. In: Analytical Biochemistry 417.1, pp. 25–35.

Ekiert, Damian C. et al. (Apr. 2009). “Antibody recognition of a highly conserved influenza virus epitope”. eng. In: Science (New York, N.Y.) 324.5924, pp. 246–251.

Gorny, Miroslaw K. et al. (Feb. 2009). “Preferential use of the VH5-51 gene segment by the human immune response to code for antibodies against the V3 domain of HIV-1”. In: Molecular Immunology 46.5, pp. 917–926.

Grabar, Pierre (June 1974). ““ SELF “ AND “ NOT-SELF “ IN IMMUNOLOGY”. In: The Lancet. Originally published as Volume 1, Issue 7870 303.7870, pp. 1320–1322.

Hecker, Michael et al. (Apr. 2016). “High-Density Peptide Microarray Analysis of IgG Autoantibody Reactivities in Serum and Cerebrospinal Fluid of Multiple Sclerosis Patients”. In: Molecular & Cellular Proteomics: MCP 15.4, pp. 1360–1380.

Höftberger, Romana et al. (Nov. 2022). “Pathogenic autoantibodies in multiple sclerosis — from a simple idea to a complex concept”. en. In: Nature Reviews Neurology 18.11. Number: 11 Publisher: Nature Publishing Group, pp. 681–688.

Kabat, E. A., M. Glusman, and V. Knaub (May 1948). “Quantitative estimation of the albumin and gamma globulin in normal and pathologic cerebrospinal fluid by immunochemical methods”. eng. In: The American Journal of Medicine 4.5, pp. 653–662.

Katoh, Masaru (June 2013). “Functional proteomics, human genetics and cancer biology of GIPC family members”. en. In: Experimental & Molecular Medicine 45.6. Publisher: Nature Publishing Group, e26–e26.

Kim, Sang Il et al. (Jan. 2021). “Stereotypic neutralizing VH antibodies against SARS-CoV-2 spike protein receptor binding domain in patients with COVID-19 and healthy individuals”. In: Science Translational Medicine 13.578. Publisher: American Association for the Advancement of Science, eabd6990.

Klein, Jochen (Sept. 2005). “Functions and pathophysiological roles of phospholipase D in the brain”. In: Journal of Neurochemistry 94.6. Publisher: John Wiley & Sons, Ltd, pp. 1473–1487.

Kornau, Hans-Christian et al. (Mar. 2020). “Human Cerebrospinal Fluid Monoclonal LGI1 Autoantibodies Increase Neuronal Excitability”. eng. In: Annals of Neurology 87.3, pp. 405–418.

Kovalenko, Oleg V., Xiuwei H. Yang, and Martin E. Hemler (Nov. 2007). “A Novel Cysteine Crosslinking Method Reveals a Direct Association between Claudin-1 and Tetraspanin CD9*”. In: Molecular & Cellular Proteomics 6.11, pp. 1855– 1867.

Kreye, Jakob et al. (2016). “Human cerebrospinal fluid monoclonal N -methyl-D-aspartate receptor autoantibodies are sufficient for encephalitis pathogenesis”. In: Brain 139.10, pp. 2641–2652.

Lanz, Tobias V. et al. (Mar. 2022). “Clonally expanded B cells in multiple sclerosis bind EBV EBNA1 and GlialCAM”. en. In: Nature 603.7900. Number: 7900 Publisher: Nature Publishing Group, pp. 321–327.

Lindeman, Ida et al. (2022). “Stereotyped B-cell responses are linked to IgG constant region polymorphisms in multiple sclerosis”. en. In: European Journal of Immunology 52.4, pp. 550–565.

McKeon, Andrew et al. (Sept. 2023). “Utility of Protein Microarrays for Detection of Classified and Novel Antibodies in Autoimmune Neurologic Disease”. In: Neurology Neuroimmunology & Neuroinflammation 10.5. Publisher: Wolters Kluwer, e200145.

Mellacheruvu, Dattatreya et al. (Aug. 2013). “The CRAPome: a contaminant repository for affinity purification–mass spectrometry data”. en. In: Nature Methods 10.8. Publisher: Nature Publishing Group, pp. 730–736.

Obermeier, Birgit et al. (June 2008). “Matching of oligoclonal immunoglobulin transcriptomes and proteomes of cerebrospinal fluid in multiple sclerosis”. en. In: Nature Medicine 14.6. Publisher: Nature Publishing Group, pp. 688–693.

Owens, Gregory P. et al. (Dec. 2011). “Viruses and Multiple Sclerosis”. en. In: The Neuroscientist 17.6, pp. 659–676.

Owens Gregory P. et al. (Dec. 15, 2006). “Screening Random Peptide Libraries with Subacute Sclerosing PanencephalitisBrain-Derived Recombinant Antibodies Identifies Multiple Epitopes in the C-Terminal Region of the Measles Virus Nucleocapsid Protein”. In: Journal of Virology 80.24. Publisher: American Society for Microbiology, pp. 12121–12130.

Polak, Justyna et al. (2023). “Single-cell transcriptomics combined with proteomics of intrathecal IgG reveal transcriptional heterogeneity of oligoclonal IgG-secreting cells in multiple sclerosis”. In: Frontiers in Cellular Neuroscience 17.

Puentes, Fabiola et al. (2017). “Neurofilament light as an immune target for pathogenic antibodies”. en. In: Immunology 152.4, pp. 580–588.

Quintana, Francisco J. et al. (Dec. 2008). “Antigen microarrays identify unique serum autoantibody signatures in clinical and pathologic subtypes of multiple sclerosis”. eng. In: Proceedings of the National Academy of Sciences of the United States of America 105.48, pp. 18889–18894.

Rand, K. H. et al. (July 1998). “Molecular approach to find target(s) for oligoclonal bands in multiple sclerosis”. eng. In: Journal of Neurology, Neurosurgery, and Psychiatry 65.1, pp. 48–55.

Ruprecht, Klemens et al. (July 2014). “Multiple sclerosis: The elevated antibody response to Epstein–Barr virus primarily targets, but is not confined to, the glycine–alanine repeat of Epstein–Barr nuclear antigen-1”. In: Journal of Neuroimmunology 272.1, pp. 56–61.

Sargsyan, S.A. et al. (Apr. 2010). “Absence of Epstein-Barr virus in the brain and CSF of patients with multiple sclerosis”. In: Neurology 74.14. Publisher: Wolters Kluwer, pp. 1127–1135.

Sathiyamoorthy, Karthik et al. (Dec. 2016). “Structural basis for Epstein–Barr virus host cell tropism mediated by gp42 and gHgL entry glycoproteins”. en. In: Nature Communications 7.1. Publisher: Nature Publishing Group, p. 13557.

Sato, Shigeo et al. (Dec. 2016). “Role for the MED21-MED7 Hinge in Assembly of the Mediator-RNA Polymerase II Holoenzyme *”. English. In: Journal of Biological Chemistry 291.52. Publisher: Elsevier, pp. 26886–26898.

Sattarnezhad, Neda et al. (Mar. 2025). “Antibody reactivity against EBNA1 and GlialCAM differentiates multiple sclerosis patients from healthy controls”. In: Proceedings of the National Academy of Sciences 122.11. Publisher: Proceedings of the National Academy of Sciences, e2424986122.

Thacker, Evan L., Fariba Mirzaei, and Alberto Ascherio (2006). “Infectious mononucleosis and risk for multiple sclerosis: A meta-analysis”. en. In: Annals of Neurology 59.3, pp. 499–503.

Theorell, Jakob et al. (Feb. 2024). “Ultrahigh frequencies of peripherally matured LGI1- and CASPR2-reactive B cells characterize the cerebrospinal fluid in autoimmune encephalitis”. en. In: Proceedings of the National Academy of Sciences 121.7, e2311049121.

Thomas, Olivia G. et al. (May 2023). “Cross-reactive EBNA1 immunity targets alpha-crystallin B and is associated with multiple sclerosis”. In: Science Advances 9.20. Publisher: American Association for the Advancement of Science, eadg3032.

Thul, Peter J. et al. (May 2017). “A subcellular map of the human proteome”. In: Science 356.6340. Publisher: American Association for the Advancement of Science, eaal3321.

Tsai, Kuang-Lei et al. (Apr. 2017). “Mediator structure and rearrangements required for holoenzyme formation”. en. In: Nature 544.7649. Publisher: Nature Publishing Group, pp. 196–201.

Tyanova, Stefka et al. (Sept. 2016). “The Perseus computational platform for comprehensive analysis of (prote)omics data”. en. In: Nature Methods 13.9. Publisher: Nature Publishing Group, pp. 731–740.

Vandvik, B. et al. (1976). “Oligoclonal Measles Virus-Specific IgG Antibodies Isolated from Cerebrospinal Fluids, Brain Extracts, and Sera from Patients with Subacute Sclerosing Panencephalitis and Multiple Sclerosis”. en. In: Scandinavian Journal of Immunology 5.8, pp. 979–992.

