## Supplementary Information for "Putative target antigens of the stereotyped intrathecal B cell response in multiple sclerosis"

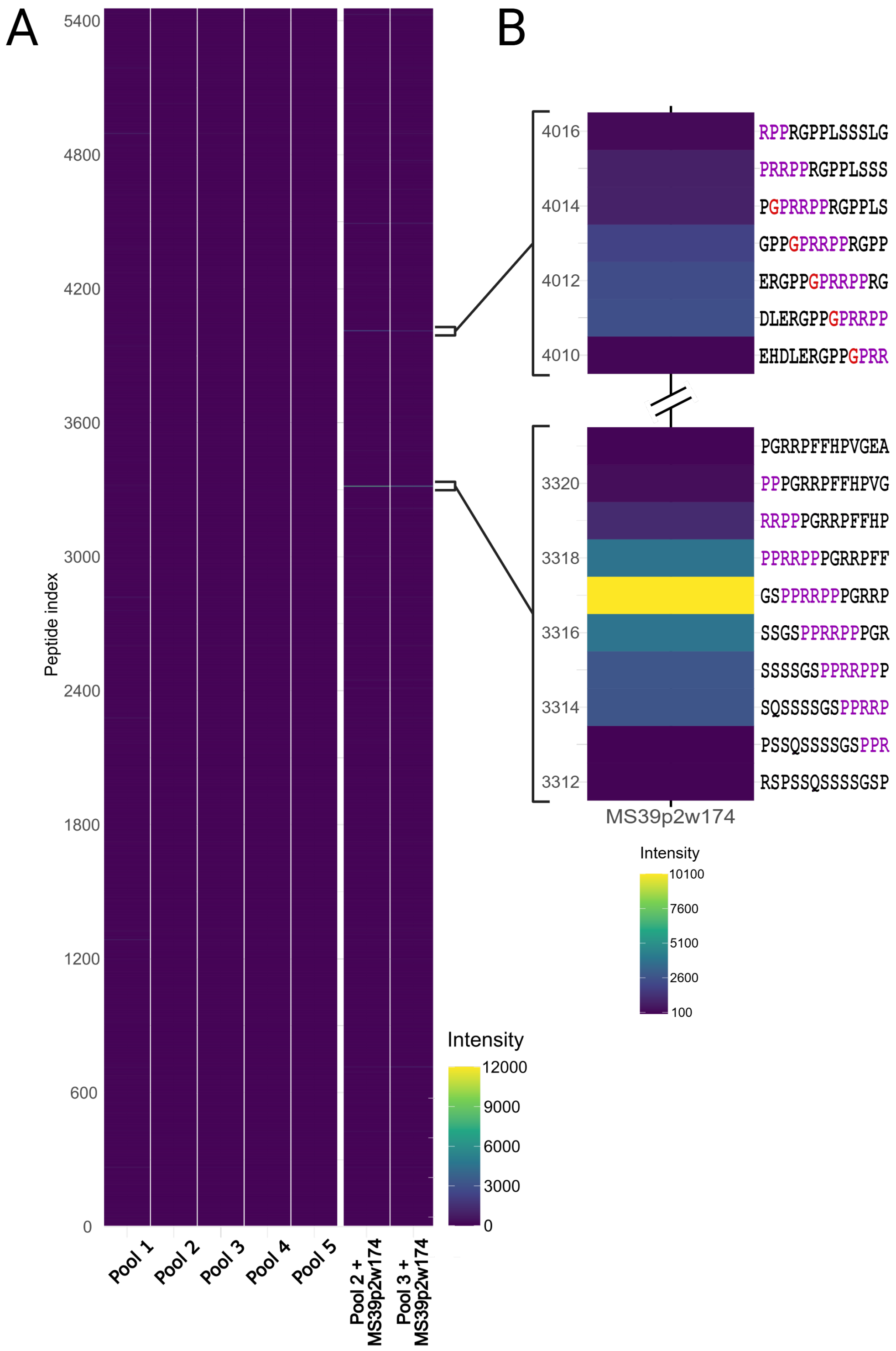
A
B
5400
4800
4200
3600
3000
2400
1800
1200
600
0
Peptide index
Pool 1
Pool 2
Pool 3
Pool 4
Pool 5
Pool 2 + MS39p2w174
Pool 3 + MS39p2w174
Intensity
12000
9000
6000
3000
0
4016
4014
4012
4010
RPPRGPPLSSSLG
PRRPPRGPPLSSS
PGPRRPPRGPPLS
GPPGPRRPPRGPP
ERGPPGPRRPPRG
DLERGPPGPRRPP
EHDLERGPPGPRR
3320
3318
3316
3314
3312
PGRRPFFHPVGEA
PPPGRRPFFHPVG
RRPPPGRRPFFHP
PPRRPPPGRRPFF
GSPPRRPPPGRRP
SSGSPPRRPPPGR
SSSSGSPPRRPPP
SQSSSSGSPPRRP
PSSQSSSSGSPPR
RSPSSQSSSSGSP
MS39p2w174
Intensity
10100
7600
5100
2600
100

**Figure S1**

**Figure S1. MS mAbs show no reactivity towards linear EBV peptides.**
(A) MS mAbs were screened against a microarray containing ca. 5,500 indexed 13-mer peptides spanning over 18 EBV proteins (y-axis). The antibodies were screened in five pools containing four to five mAbs per pool (x-axis). The chips that were used to screen Pool 2 and Pool 3 were subsequently incubated with MS39p2w179 (x-axis, two last panels). The resulting verification of the previously published epitope of this antibody is shown in the bottom panel of (B), where the published epitope is marked in purple. A second signal, showing a peptide with very similar sequence to the published epitope, is shown in the top panel. The deviating amino acid is marked red. All intensity values are given as uncorrected arbitrary units (AU). Panel (B) shows the mean intensity value of two independent experiments.

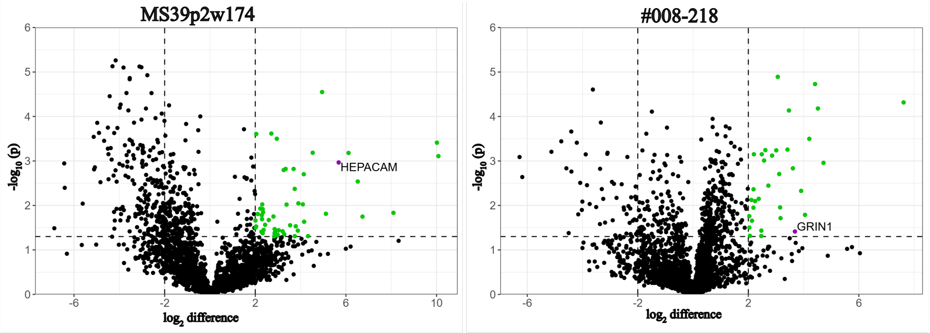


**Figure S2. Results from IP-MS/MS performed on of human brain lysate for the control antibodies.** The known target of the antibody is named and marked purple.

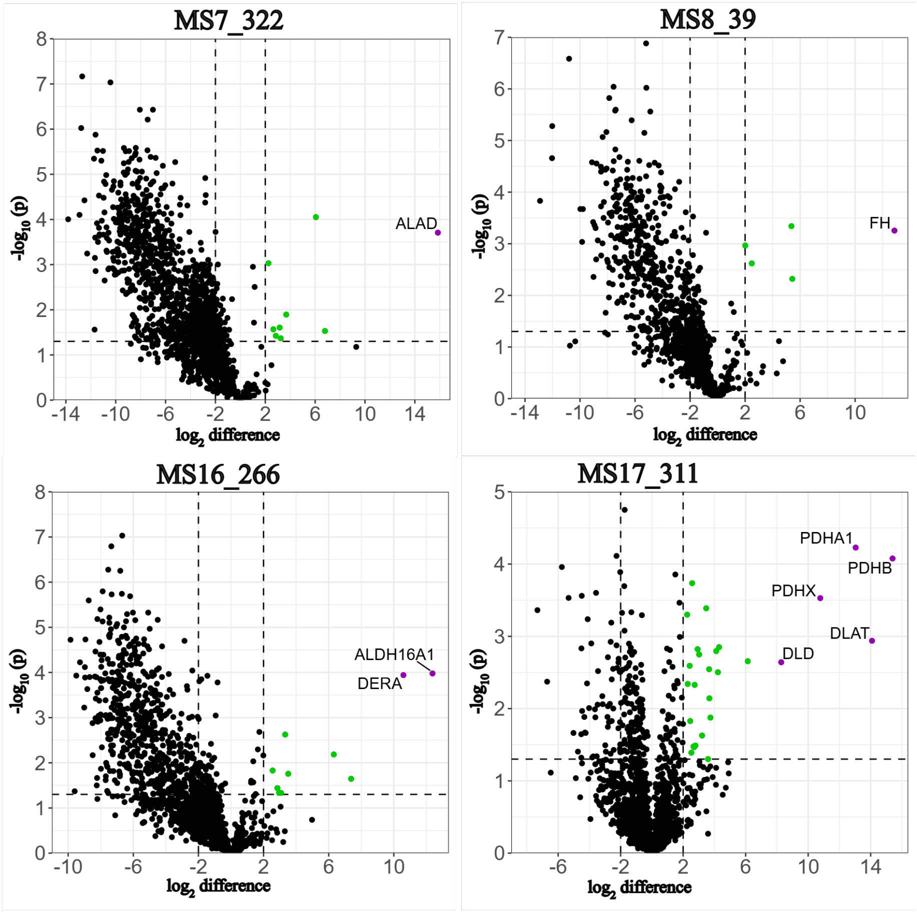


**Figure S3. Examples of MS mAbs that clearly enriched one protein or several proteins that are known to be in direct contact in the cell.** None of these proteins were enriched by other mAbs.

**Table S1.** Putative antigenic targets from HuProt tested on ELISA

| Protein used for ELISA | cat# | Producer |
|---|---|---|
| MED21 | PRO-1476 | ProSpec (Rehovot, Israel) |
| GIPC2 | PRO-1035 | ProSpec (Rehovot, Israel) |
| KHDRBS2 | TP305428 | OriGene (Rockville, MD, USA) |
| SCL-70 (=TOP1) | 17455-H07B | SinoBiological (Beijing, China) |
| NAPB | Ag5239 | Proteintech (Manchester, UK) |
| SYTL1 | CSB-MP815582HU | Cusabio (Houston, TX, USA) |
| PPIA | ab290069 | Abcam (Cambridge, UK) |

**Table S2.** Summary of quality control experiments of MS mAbs. Slight deviances from the norm in some experiments prompted the addition of extra experiments for some antibodies (last two columns). NA indicates that the experiment was not performed for the indicated antibody.

| **Antibody ID** | **SignalP 6.0 in-silico SS cleavage prediticion** | **Reducing SDS-PAGE** | **Nonreducing SDS-PAGE** | **SEC curve indicates monomer** | **Plasmid re-sequencing** | **Mass spectrometry V-region peptides** |
|---|---|---|---|---|---|---|
| MS1_300 | PASS | †1 | PASS | PASS | PASS | NA |
| MS2_25 | PASS | PASS | PASS | PASS | NA | NA |
| MS2_30 | PASS | PASS | PASS | PASS | NA | NA |
| MS2_80 * | PASS | PASS | †5 | PASS | PASS | PASS |
| MS6_274 | PASS | PASS | PASS | PASS | NA | NA |
| MS7_307 | PASS | PASS | PASS | PASS | PASS | NA |
| MS7_322 | PASS | PASS | PASS | PASS | NA | NA |
| MS7_372 | PASS | †2 | †5, 6, 7 | PASS | PASS | PASS |
| MS8_39 | PASS | PASS | PASS | PASS | NA | NA |
| MS8_78 | PASS | †1 | PASS | PASS | PASS | NA |
| MS8_274 | PASS | PASS | PASS | PASS | NA | NA |
| MS10_88 | PASS | PASS | PASS | PASS†8 | PASS | PASS |
| MS10_146 | PASS | PASS | PASS | PASS | NA | NA |
| MS13_50 | PASS | †2, 3 | †5 | PASS | PASS | NA |
| MS13_58 | PASS | PASS | PASS | PASS | PASS | NA |
| MS13_73 | PASS | PASS | PASS | PASS | NA | NA |
| MS14_162 | PASS | PASS | PASS | NA | NA | NA |
| MS14_165 | PASS | PASS | PASS | PASS | NA | NA |
| MS16_266 | PASS | PASS | †5, 6 | PASS | PASS | PASS |
| MS17_311 | PASS | †3 | PASS | PASS | PASS | NA |
| MS19_38 | PASS | †1 | PASS | PASS | PASS | NA |
| MS19_85 | PASS | PASS | PASS | PASS | NA | NA |
| MS20_14 | PASS | PASS | †5, 6 | PASS | PASS | PASS |
| MS21_115 | PASS | PASS | PASS | PASS | PASS | NA |
| MS39p2w174 ** | PASS | PASS | PASS | PASS | NA | NA |
| #008-218 ** | PASS | †4 | PASS | PASS | NA | NA |
| MS5_NB2 | PASS | PASS | PASS | PASS | NA | NA |

**Shown in example figure. **Bound to target.*

*† **1** Double H band, **2** H chain aggregates, **3** Large L band, **4** Diffuse L band, **5** Appears smaller, **6** diffuse band, **7** no H chain aggreg., **8** appeared with elongated tail*

**Table S3.** Overview of mAbs and proteins enriched in IP-MS/MS. Antibodies that enriched the protein in question are indicated with a "1" in their respective column.

| Gene name | MS14_165 | MS21_115 | MS8_178 | MS13_73 | MS2_80 | MS7_372 | MS8_39 | MS13_58 | MS7_322 | MS7_307 | MS19_85 | MS16_266 | MS6_274 | MS2_30 | MS2_25 | MS17_311 | MS19_38 | MS1_300 | MS10_146 | MS14_162 | MS13_50 | MS10_88 | MS20_14 | MS8_78 | Sum |
|---|---|---|---|---|---|---|---|---|---|---|---|---|---|---|---|---|---|---|---|---|---|---|---|---|---|
| CLDN11 | 0 | 0 | 0 | 0 | 0 | 0 | 0 | 0 | 0 | 0 | 0 | 0 | 0 | 0 | 0 | 1 | 1 | 0 | 1 | 0 | 0 | 1 | 1 | 0 | 5 |
| ANK3 | 0 | 0 | 0 | 0 | 0 | 0 | 0 | 0 | 0 | 0 | 0 | 0 | 0 | 0 | 0 | 0 | 0 | 0 | 0 | 0 | 1 | 1 | 1 | 1 | 4 |
| CD9 | 0 | 0 | 0 | 0 | 0 | 0 | 0 | 0 | 0 | 0 | 0 | 0 | 0 | 0 | 0 | 0 | 1 | 0 | 1 | 0 | 0 | 1 | 1 | 0 | 4 |
| OPALIN | 0 | 0 | 0 | 0 | 0 | 0 | 0 | 0 | 0 | 0 | 0 | 0 | 0 | 0 | 0 | 0 | 0 | 0 | 1 | 0 | 1 | 1 | 0 | 0 | 3 |
| S100B | 0 | 0 | 0 | 0 | 0 | 0 | 0 | 0 | 0 | 0 | 1 | 0 | 0 | 0 | 0 | 0 | 1 | 0 | 0 | 1 | 0 | 0 | 0 | 0 | 3 |
| PLD1 | 0 | 0 | 0 | 0 | 0 | 0 | 1 | 1 | 1 | 0 | 1 | 0 | 0 | 1 | 1 | 0 | 0 | 0 | 0 | 0 | 0 | 0 | 0 | 0 | 6 |
| CORO2B | 0 | 0 | 0 | 0 | 0 | 0 | 0 | 0 | 0 | 0 | 0 | 0 | 0 | 0 | 0 | 0 | 0 | 0 | 0 | 1 | 1 | 1 | 1 | 1 | 5 |
| ANK2 | 0 | 0 | 0 | 0 | 0 | 0 | 0 | 0 | 0 | 0 | 0 | 0 | 0 | 0 | 0 | 0 | 0 | 0 | 0 | 1 | 1 | 1 | 1 | 1 | 5 |
| CAMK2G | 0 | 0 | 0 | 0 | 0 | 0 | 0 | 0 | 0 | 0 | 0 | 0 | 0 | 0 | 0 | 0 | 0 | 0 | 0 | 1 | 1 | 1 | 1 | 1 | 5 |
| FBXO2 | 0 | 0 | 0 | 0 | 0 | 0 | 0 | 0 | 0 | 0 | 0 | 0 | 0 | 0 | 0 | 1 | 0 | 0 | 0 | 1 | 1 | 0 | 1 | 1 | 5 |
| PRKCG | 0 | 0 | 0 | 0 | 0 | 0 | 0 | 0 | 0 | 0 | 0 | 0 | 0 | 0 | 0 | 0 | 0 | 0 | 0 | 1 | 1 | 1 | 1 | 1 | 5 |
| SNTA1 | 0 | 0 | 0 | 0 | 0 | 0 | 0 | 0 | 0 | 0 | 0 | 0 | 0 | 0 | 0 | 0 | 0 | 0 | 0 | 0 | 1 | 1 | 1 | 1 | 4 |
| PALM | 0 | 0 | 0 | 0 | 0 | 0 | 0 | 0 | 0 | 0 | 0 | 0 | 0 | 0 | 0 | 0 | 0 | 0 | 0 | 0 | 1 | 1 | 1 | 1 | 4 |
| GLS | 0 | 0 | 0 | 0 | 0 | 0 | 0 | 0 | 0 | 0 | 0 | 0 | 0 | 0 | 0 | 1 | 0 | 0 | 1 | 0 | 0 | 0 | 1 | 1 | 4 |
| DNM3 | 0 | 0 | 0 | 0 | 0 | 0 | 0 | 0 | 0 | 0 | 0 | 0 | 0 | 0 | 0 | 0 | 0 | 0 | 1 | 0 | 1 | 0 | 1 | 1 | 4 |
| GNAI1 | 0 | 0 | 0 | 0 | 0 | 0 | 0 | 0 | 0 | 0 | 0 | 0 | 0 | 0 | 0 | 0 | 1 | 0 | 1 | 0 | 1 | 1 | 0 | 0 | 4 |
| MAPRE2 | 0 | 0 | 0 | 0 | 0 | 0 | 0 | 0 | 0 | 0 | 0 | 0 | 0 | 0 | 0 | 0 | 0 | 0 | 1 | 1 | 0 | 0 | 1 | 0 | 3 |
| PSD3 | 0 | 0 | 0 | 0 | 0 | 0 | 0 | 0 | 0 | 0 | 0 | 0 | 0 | 0 | 0 | 0 | 0 | 0 | 0 | 0 | 1 | 1 | 0 | 1 | 3 |
| CTNNA2 | 0 | 0 | 0 | 0 | 0 | 0 | 0 | 0 | 0 | 0 | 0 | 1 | 0 | 0 | 0 | 0 | 0 | 0 | 0 | 0 | 0 | 0 | 1 | 1 | 3 |
| CAMKV | 0 | 0 | 0 | 0 | 0 | 0 | 0 | 0 | 0 | 0 | 0 | 0 | 0 | 0 | 0 | 0 | 0 | 0 | 0 | 1 | 0 | 0 | 1 | 1 | 3 |
| SLC12A5 | 0 | 0 | 0 | 0 | 0 | 0 | 0 | 0 | 0 | 0 | 0 | 0 | 0 | 0 | 0 | 0 | 0 | 0 | 0 | 0 | 0 | 1 | 1 | 1 | 3 |
| DMTN | 0 | 0 | 0 | 0 | 0 | 0 | 0 | 0 | 0 | 0 | 0 | 0 | 0 | 0 | 0 | 0 | 0 | 0 | 0 | 0 | 0 | 1 | 1 | 1 | 3 |
| TJP2 | 0 | 0 | 0 | 0 | 0 | 0 | 0 | 0 | 1 | 0 | 0 | 1 | 0 | 0 | 1 | 0 | 0 | 0 | 0 | 0 | 0 | 0 | 0 | 0 | 3 |
| PLEKHB1 | 0 | 0 | 0 | 0 | 0 | 0 | 0 | 0 | 0 | 0 | 0 | 0 | 0 | 0 | 0 | 0 | 0 | 0 | 1 | 0 | 1 | 1 | 0 | 0 | 3 |
| PHF24 | 0 | 0 | 0 | 0 | 0 | 0 | 0 | 0 | 0 | 0 | 0 | 0 | 0 | 0 | 0 | 0 | 0 | 0 | 0 | 0 | 0 | 1 | 1 | 1 | 3 |
| PRKCB | 0 | 0 | 0 | 0 | 0 | 0 | 0 | 0 | 0 | 0 | 0 | 0 | 0 | 0 | 0 | 0 | 0 | 0 | 0 | 1 | 0 | 0 | 1 | 1 | 3 |
| RAB24 | 0 | 0 | 0 | 0 | 0 | 0 | 0 | 0 | 0 | 0 | 0 | 0 | 0 | 0 | 0 | 0 | 0 | 0 | 0 | 0 | 1 | 1 | 0 | 1 | 3 |
| MYO5A | 0 | 0 | 0 | 0 | 1 | 1 | 0 | 0 | 0 | 1 | 1 | 0 | 0 | 0 | 0 | 0 | 0 | 0 | 0 | 1 | 1 | 1 | 1 | 1 | 9 |
| MYH10 | 0 | 0 | 0 | 0 | 0 | 0 | 0 | 0 | 0 | 0 | 0 | 0 | 0 | 1 | 0 | 0 | 0 | 1 | 0 | 1 | 1 | 1 | 1 | 1 | 7 |
| GSN | 0 | 0 | 0 | 0 | 0 | 0 | 0 | 0 | 0 | 0 | 0 | 0 | 0 | 0 | 0 | 0 | 0 | 1 | 1 | 1 | 1 | 1 | 1 | 1 | 7 |
| MYO1D | 0 | 0 | 0 | 0 | 0 | 0 | 0 | 0 | 0 | 0 | 0 | 0 | 0 | 0 | 0 | 0 | 1 | 1 | 1 | 0 | 1 | 1 | 1 | 1 | 7 |
| TMOD2 | 0 | 0 | 0 | 0 | 0 | 0 | 0 | 0 | 0 | 0 | 0 | 0 | 0 | 0 | 0 | 0 | 0 | 1 | 0 | 1 | 1 | 1 | 1 | 1 | 6 |
| SPTAN1 | 0 | 0 | 0 | 0 | 0 | 0 | 0 | 0 | 0 | 0 | 0 | 0 | 0 | 0 | 0 | 0 | 0 | 1 | 0 | 1 | 1 | 1 | 1 | 1 | 6 |
| ACTN1 | 0 | 0 | 0 | 0 | 0 | 0 | 0 | 1 | 0 | 0 | 0 | 0 | 0 | 0 | 0 | 0 | 0 | 0 | 0 | 1 | 1 | 1 | 1 | 1 | 6 |
| KIF21A | 0 | 0 | 0 | 0 | 0 | 0 | 0 | 1 | 0 | 0 | 1 | 0 | 0 | 1 | 0 | 0 | 0 | 0 | 0 | 1 | 0 | 0 | 1 | 1 | 6 |
| MYH9 | 0 | 0 | 0 | 0 | 0 | 0 | 0 | 0 | 0 | 0 | 0 | 0 | 0 | 0 | 0 | 0 | 0 | 1 | 0 | 1 | 1 | 1 | 1 | 1 | 6 |
| SPTBN1 | 0 | 0 | 0 | 0 | 0 | 0 | 0 | 0 | 0 | 0 | 0 | 0 | 0 | 0 | 0 | 0 | 0 | 0 | 0 | 1 | 1 | 1 | 1 | 1 | 5 |
| DBN1 | 0 | 0 | 0 | 0 | 0 | 0 | 0 | 0 | 0 | 0 | 0 | 0 | 0 | 0 | 0 | 0 | 0 | 0 | 0 | 1 | 1 | 1 | 1 | 1 | 5 |
| CAMK2D | 0 | 0 | 0 | 0 | 0 | 0 | 0 | 0 | 0 | 0 | 0 | 0 | 0 | 0 | 0 | 0 | 0 | 0 | 0 | 1 | 1 | 1 | 1 | 1 | 5 |
| VDAC2 | 0 | 0 | 0 | 0 | 0 | 0 | 0 | 0 | 1 | 0 | 0 | 0 | 1 | 0 | 0 | 0 | 0 | 0 | 1 | 0 | 0 | 0 | 1 | 1 | 5 |
| CAP2 | 0 | 0 | 0 | 0 | 0 | 0 | 0 | 0 | 0 | 0 | 0 | 0 | 0 | 0 | 0 | 0 | 0 | 0 | 0 | 1 | 1 | 1 | 1 | 1 | 5 |
| ABLIM1 | 0 | 0 | 0 | 0 | 0 | 0 | 0 | 0 | 1 | 0 | 0 | 0 | 0 | 0 | 1 | 0 | 0 | 0 | 0 | 0 | 1 | 1 | 0 | 1 | 5 |
| CALM3;CALM2;CALM1 | 0 | 0 | 0 | 0 | 0 | 0 | 0 | 0 | 0 | 0 | 0 | 0 | 0 | 0 | 0 | 0 | 0 | 0 | 0 | 1 | 1 | 1 | 1 | 1 | 5 |
| TPM3 | 0 | 0 | 0 | 0 | 0 | 0 | 0 | 0 | 0 | 1 | 0 | 0 | 0 | 0 | 0 | 0 | 0 | 1 | 0 | 0 | 1 | 1 | 0 | 1 | 5 |
| LIMCH1 | 0 | 0 | 0 | 0 | 0 | 0 | 0 | 0 | 0 | 1 | 0 | 1 | 0 | 0 | 1 | 0 | 0 | 1 | 0 | 0 | 0 | 0 | 1 | 0 | 5 |
| MYH14 | 0 | 0 | 0 | 0 | 0 | 0 | 0 | 0 | 0 | 0 | 0 | 0 | 0 | 0 | 0 | 0 | 0 | 1 | 0 | 0 | 1 | 1 | 1 | 1 | 5 |
| ANXA1 | 0 | 0 | 0 | 0 | 0 | 0 | 0 | 0 | 0 | 0 | 0 | 0 | 0 | 0 | 0 | 0 | 0 | 0 | 0 | 1 | 1 | 1 | 1 | 1 | 5 |
| LLGL1 | 0 | 0 | 0 | 0 | 0 | 0 | 0 | 0 | 0 | 0 | 0 | 0 | 0 | 0 | 0 | 0 | 1 | 0 | 0 | 0 | 1 | 0 | 1 | 1 | 4 |
| ADD3 | 0 | 0 | 0 | 0 | 0 | 0 | 0 | 0 | 0 | 0 | 0 | 0 | 0 | 0 | 0 | 0 | 0 | 0 | 0 | 0 | 1 | 1 | 1 | 1 | 4 |
| SPTBN2 | 0 | 0 | 0 | 0 | 0 | 0 | 0 | 0 | 0 | 0 | 0 | 0 | 0 | 0 | 0 | 0 | 0 | 0 | 0 | 0 | 1 | 1 | 1 | 1 | 4 |
| CTNNB1 | 0 | 0 | 0 | 0 | 0 | 0 | 0 | 0 | 0 | 0 | 0 | 0 | 0 | 0 | 0 | 0 | 0 | 0 | 0 | 0 | 1 | 1 | 1 | 1 | 4 |
| VDAC1 | 0 | 0 | 0 | 0 | 0 | 0 | 0 | 0 | 0 | 0 | 0 | 0 | 1 | 0 | 0 | 0 | 0 | 0 | 1 | 0 | 0 | 0 | 1 | 1 | 4 |
| MACF1 | 0 | 0 | 0 | 0 | 0 | 0 | 0 | 0 | 0 | 0 | 0 | 0 | 0 | 0 | 0 | 0 | 0 | 0 | 0 | 0 | 1 | 1 | 1 | 1 | 4 |
| IMMT | 0 | 0 | 0 | 0 | 0 | 0 | 0 | 0 | 0 | 0 | 0 | 0 | 0 | 1 | 0 | 0 | 0 | 0 | 1 | 0 | 0 | 0 | 1 | 1 | 4 |

**Table S3.** – continued from previous page

| Gene name | MS14_165 | MS21_115 | MS8_178 | MS13_73 | MS2_80 | MS7_372 | MS8_39 | MS13_58 | MS7_322 | MS7_307 | MS19_85 | MS16_266 | MS6_274 | MS2_30 | MS2_25 | MS17_311 | MS19_38 | MS1_300 | MS10_146 | MS14_162 | MS13_50 | MS10_88 | MS20_14 | MS8_78 | **Sum** |
|---|---|---|---|---|---|---|---|---|---|---|---|---|---|---|---|---|---|---|---|---|---|---|---|---|---|
| MGST3 | 0 | 0 | 0 | 0 | 1 | 0 | 0 | 0 | 0 | 0 | 0 | 1 | 1 | 0 | 0 | 0 | 0 | 0 | 0 | 0 | 0 | 0 | 1 | 0 | 4 |
| EFHD2 | 0 | 0 | 0 | 0 | 0 | 0 | 0 | 0 | 0 | 0 | 0 | 0 | 0 | 0 | 0 | 0 | 0 | 0 | 0 | 0 | 1 | 1 | 1 | 1 | 4 |
| CAPZA2 | 0 | 0 | 0 | 0 | 0 | 0 | 0 | 0 | 0 | 0 | 0 | 0 | 0 | 0 | 0 | 0 | 0 | 0 | 0 | 1 | 1 | 1 | 0 | 1 | 4 |
| PLEC | 0 | 0 | 0 | 0 | 0 | 0 | 0 | 0 | 0 | 0 | 0 | 0 | 0 | 0 | 0 | 0 | 0 | 0 | 0 | 0 | 1 | 1 | 1 | 1 | 4 |
| TPM4 | 0 | 0 | 0 | 0 | 0 | 0 | 0 | 0 | 0 | 0 | 0 | 0 | 0 | 0 | 0 | 0 | 0 | 0 | 0 | 0 | 1 | 1 | 1 | 1 | 4 |
| SYNPO | 0 | 0 | 0 | 0 | 0 | 0 | 0 | 0 | 0 | 0 | 0 | 0 | 0 | 0 | 0 | 0 | 0 | 0 | 0 | 0 | 1 | 1 | 1 | 1 | 4 |
| HSPB1 | 0 | 0 | 0 | 0 | 0 | 0 | 0 | 0 | 0 | 0 | 0 | 0 | 0 | 0 | 0 | 0 | 0 | 0 | 0 | 1 | 1 | 1 | 0 | 1 | 4 |
| MYO1C | 0 | 0 | 0 | 0 | 0 | 0 | 0 | 0 | 0 | 0 | 0 | 0 | 0 | 0 | 0 | 0 | 0 | 0 | 1 | 0 | 1 | 1 | 0 | 1 | 4 |
| VDAC3 | 0 | 0 | 0 | 0 | 0 | 0 | 0 | 0 | 0 | 0 | 0 | 0 | 1 | 0 | 0 | 0 | 0 | 0 | 1 | 0 | 0 | 0 | 1 | 1 | 4 |
| TPM1 | 0 | 0 | 0 | 0 | 0 | 0 | 0 | 0 | 0 | 0 | 0 | 0 | 0 | 0 | 0 | 0 | 0 | 1 | 0 | 0 | 1 | 1 | 0 | 1 | 4 |
| EPB41L1 | 0 | 0 | 0 | 0 | 0 | 0 | 0 | 0 | 0 | 0 | 0 | 0 | 0 | 0 | 0 | 0 | 0 | 0 | 1 | 0 | 0 | 0 | 1 | 1 | 3 |
| PIN1 | 0 | 0 | 0 | 0 | 0 | 0 | 0 | 0 | 0 | 0 | 0 | 0 | 0 | 0 | 0 | 0 | 0 | 0 | 1 | 1 | 0 | 0 | 1 | 0 | 3 |
| MYO6 | 0 | 0 | 0 | 0 | 0 | 0 | 0 | 0 | 0 | 0 | 0 | 0 | 0 | 0 | 0 | 0 | 0 | 0 | 0 | 0 | 1 | 1 | 0 | 1 | 3 |
| HK1 | 0 | 0 | 0 | 0 | 0 | 0 | 0 | 0 | 0 | 0 | 0 | 0 | 0 | 0 | 0 | 0 | 0 | 0 | 1 | 0 | 0 | 0 | 1 | 1 | 3 |
| PHB1 | 0 | 0 | 0 | 0 | 0 | 0 | 0 | 0 | 0 | 0 | 0 | 1 | 0 | 0 | 0 | 0 | 0 | 0 | 1 | 0 | 0 | 1 | 0 | 0 | 3 |
| MYO18A | 0 | 0 | 0 | 0 | 0 | 0 | 0 | 0 | 0 | 0 | 0 | 0 | 0 | 0 | 0 | 0 | 0 | 0 | 0 | 0 | 1 | 1 | 0 | 1 | 3 |
| CHCHD3 | 0 | 0 | 0 | 0 | 0 | 0 | 0 | 0 | 0 | 0 | 0 | 0 | 0 | 0 | 0 | 1 | 0 | 0 | 1 | 0 | 0 | 0 | 1 | 0 | 3 |
| SH3BGRL2 | 0 | 0 | 0 | 0 | 0 | 0 | 0 | 0 | 0 | 0 | 0 | 0 | 0 | 0 | 0 | 0 | 0 | 0 | 0 | 0 | 1 | 1 | 0 | 1 | 3 |
| SIRT2 | 0 | 0 | 0 | 0 | 0 | 0 | 0 | 0 | 0 | 0 | 0 | 0 | 0 | 0 | 0 | 0 | 1 | 0 | 0 | 0 | 1 | 1 | 0 | 0 | 3 |
| ADD1 | 0 | 0 | 0 | 0 | 0 | 0 | 0 | 0 | 0 | 0 | 0 | 0 | 0 | 0 | 0 | 0 | 0 | 0 | 0 | 0 | 0 | 1 | 1 | 1 | 3 |
| UBXN6 | 0 | 0 | 0 | 0 | 0 | 0 | 0 | 0 | 0 | 0 | 0 | 0 | 0 | 0 | 0 | 0 | 0 | 0 | 1 | 0 | 1 | 0 | 0 | 1 | 3 |
| ERP44 | 0 | 0 | 0 | 0 | 0 | 0 | 0 | 0 | 0 | 0 | 0 | 1 | 0 | 0 | 0 | 1 | 0 | 0 | 0 | 1 | 0 | 0 | 0 | 0 | 3 |
| PHB2 | 0 | 0 | 0 | 0 | 0 | 0 | 0 | 0 | 0 | 0 | 0 | 1 | 0 | 0 | 0 | 0 | 0 | 0 | 1 | 0 | 0 | 1 | 0 | 0 | 3 |
| SORBS1 | 0 | 0 | 0 | 0 | 0 | 0 | 0 | 0 | 0 | 0 | 0 | 0 | 0 | 0 | 0 | 0 | 0 | 0 | 0 | 0 | 1 | 1 | 0 | 1 | 3 |
| ACTN2 | 0 | 0 | 0 | 0 | 0 | 0 | 0 | 0 | 0 | 0 | 0 | 0 | 0 | 0 | 0 | 0 | 0 | 0 | 0 | 0 | 1 | 1 | 0 | 1 | 3 |
| DST | 0 | 0 | 0 | 1 | 0 | 0 | 0 | 0 | 0 | 0 | 0 | 0 | 0 | 0 | 0 | 1 | 0 | 0 | 0 | 0 | 0 | 0 | 0 | 1 | 3 |
| MYL6 | 0 | 0 | 0 | 0 | 0 | 0 | 0 | 0 | 0 | 0 | 0 | 0 | 0 | 0 | 0 | 0 | 0 | 1 | 0 | 0 | 1 | 0 | 0 | 1 | 3 |
| ACTB | 0 | 0 | 0 | 0 | 0 | 0 | 0 | 0 | 0 | 0 | 0 | 0 | 0 | 0 | 0 | 0 | 0 | 0 | 0 | 0 | 1 | 1 | 0 | 1 | 3 |
| TMOD1 | 0 | 0 | 0 | 0 | 0 | 0 | 0 | 0 | 0 | 0 | 0 | 0 | 0 | 0 | 0 | 0 | 0 | 0 | 0 | 0 | 1 | 1 | 0 | 1 | 3 |
| ACTBL2 | 0 | 0 | 0 | 0 | 0 | 0 | 0 | 0 | 0 | 0 | 0 | 0 | 0 | 0 | 0 | 0 | 0 | 0 | 1 | 0 | 0 | 1 | 0 | 1 | 3 |

**Table S4.** Summary of mAb-enriched proteins, their function and location

| Gene name | RNA (brain) | Protein (brain) | RNA (OGs)* | CRAPome score** | General localization | Function |
|---|---|---|---|---|---|---|
| CLDN11 | high | mid | high | 0 | Membranous expression in sertoli cells in testis and oligodendrocytes in the CNS | Tight junctions in CNS myelin, oligodendrocyte proliferation, and blood-testis barrier. |
| ANK3 | high | high | high | 60 | Membranous expression in several tissues. | Membrane-cytoskeleton linker. May participate in the maintenance/targeting of ion channels and cell adhesion molecules at the nodes of Ranvier and axonal initial segments |
| CD9 | mid | high | low | 0 | Ubiquitous cytoplasmic expression. | Regulates cell processes like sperm-egg fusion, platelet activation, cell adhesion, and prevents undesired cell fusion. |
| OPALIN | high | high | high | 0 | Selective expression in CNS. | Increases myelin gene expression and promotes oligodendrocyte differentiation. |
| S100B | high | high | mid | 0 | Nuclear and cytoplasmic expression mainly in CNS, peripheral nerves, adipocytes and subset of lymphoid cells. | Small zinc- and- and calcium-binding protein |
| PLD1 | low | low | high | 3 | Cytoplasmic and membranous expression at variable levels in most tissues | Function as phospholipase selective for phosphatidylcholine |
| CORO2B | high | high | high | 10 | General cytoplasmic expression, most abundant in CNS (OG). | May play a role in the reorganization of neuronal actin structure. |
| ANK2 | high | high | high | 56 | Cytoplasmic and membranous expression with high abundance in the brain, peripheral nerves, renal collecting ducts and the adrenal gland. | Plays an essential role in the localization and membrane stabilization of ion transporters and ion channels in several cell types, including cardiomyocytes, as well as in striated muscle cells. Plays a role in endocytosis and intracellular protein transport. |
| CAMK2G | high | high | mid | 34 | Cytoplasmic expression at variable levels in selected tissues, most abundant neuronal cells. | This protein kinase, after activation, regulates calcium transport in muscle and may influence neuronal structure and plasticity. |
| FBXO2 | high | mid | mid | 2 | Cytoplasmic expression in brain. | Recognizes unfolded glycoproteins for ERAD, promotes ubiquitination and degradation, prevents cytosolic aggregates |
| PRKCG | high | high | low | 16 | Selective cytoplasmic expression in Purkinje cells and neuropil. | Kinase that regulates neuronal receptors, synaptic function, pain, and stress responses. |
| SNTA1 | mid | high | low | 2 | Mainly cytoplasmic expression in most tissues. | Organizes membrane protein localization and links receptors to cytoskeleton; crucial for synapse formation. |
| PALM | high | mid | low | 11 | High expression of renal tubules and neuropil. | Involved in membrane dynamics; essential for filopodia, spine maturation, and synapse formation. |
| GLS | mid | mid | mid | 54 | High granular cytoplasmic expression in renal tubules and neuronal cells. | Catalyzes glutamine catabolism, maintains acid-base balance, regulates brain glutamate. |
| DNM3 | high | N/A | high | 56 | Intracellular expression in brain | associate with microtubules and are involved in vesicular transport. |
| GNAI1 | high | mid | high | 155 | Cytoplasmic expression in most tissues. | Mediates GPCR signaling, regulates adenylate cyclase, and essential for cytokinesis. |
| MAPRE2 | high | high | high | 101 | Cytoplasmic expression in several tissues, most abundant in the CNS | May be involved in microtubule polymerization, and spindle function by stabilizing microtubules and anchoring them at centrosomes. |

| | | | | | | |
|---|---|---|---|---|---|---|
| PSD3 | high | high | high | 143 | Highly expressed in neuronal cells. | Guanine nucleotide exchange factor for ARF6. |
| CTNNA2 | high | mid | mid | 31 | Distinct expression in neuropil. | May function as a linker between cadherin adhesion receptors and the cytoskeleton to regulate cell-cell adhesion and differentiation in the nervous system |
| CAMKV | high | mid | low | 0 | Selective expression in CNS. | Predicted to enable calmodulin binding activity and calmodulin-dependent protein kinase activity. Predicted to be involved in peptidyl-serine phosphorylation. Predicted to be located in cytoplasmic vesicle membrane and plasma membrane. Predicted to be active in glutamatergic synapse and postsynapse |
| SLC12A5 | high | mid | low | 1 | Selective cytoplasmic expression in CNS. | Mediates potassium-chloride cotransport in neurons, crucial for Cl(-) homeostasis, GABA-A and glycine receptor function, and dendritic spine formation. |
| DMTN | high | mid | low | 3 | Cytoplasmic expression in several tissue. Additional positivity in red blood cells. Tissue enhanced (Brain). | Induces F-actin bundles, stabilizes erythrocyte membrane, regulates actin dynamics |
| TJP2 | mid | low | high | 106 | Cytoplasmic and membranous expression in most tissues. | Component of tight junctions in epithelial and endothelial cells, essential for junction assembly. |
| PLEKHB1 | high | N/A | low | 0 | Mainly membranous expression in brain. | Predicted to enable protein C-terminus binding activity and protein homodimerization activity. |
| PHF24 | high | high | mid | 0 | Expression in CNS and testis. | Predicted to bind metal ions, involved in pain perception, GABA signaling, and regulation of GABAergic synaptic transmission. |
| PRKCB | high | mid | mid | 6 | Cytoplasmic and nuclear expression in lymphoid tissues and CNS. | This protein kinase has been reported to be involved in many different cellular functions, such as B cell activation, apoptosis induction, endothelial cell proliferation, and intestinal sugar absorption |
| RAB24 | high | N/A | mid | 1 | Localized to the cytosol in several tissues | a small GTPase of the Rab subfamily of Ras-related proteins that regulate intracellular protein trafficking |
| MYO5A | high | high | high | 52 | Cytoplasmic expression in several tissues, most abundant in CNS | Actin-based motor involved in melanosome and vesicle transport, aids dendrite formation. |
| MYH10 | mid | high | mid | 379 | Cytoplasmic and membranous expression in most tissues. Highest expression in Purkinje cells, trophoblastic cells and cells in renal tubules. | Involved in cytokinesis, cell shape, secretion, and type I collagen mRNA stabilization. |
| GSN | low | high | mid | 49 | Cytoplasmic expression mainly in macrophages in several tissues, and in renal glomeruli, glial cells and in islets of Langerhans | Calcium-regulated, actin-modulating protein that binds to the plus (or barbed) ends of actin monomers or filaments, preventing monomer exchange |
| MYO1D | mid | mid | high | 57 | Cytoplasmic expression in most tissues, oligodendocyte enriched. | Facilitates actin-based motor function, endosomal trafficking, and ciliary organization. |
| TMOD2 | high | high | high | 51 | Cytoplasmic expression in CNS. | Blocks actin filament dynamics, stabilizes membrane skeleton structure. |
| SPTAN1 | high | high | mid | 324 | General cytoplasmic and membranous expression in most tissues. | Interacts with calmodulin for calcium-dependent cytoskeleton movement at the membrane. |

| | | | | | | |
|---|---|---|---|---|---|---|
| ACTN1 | low | high | low | 314 | High cytoplasmic and membranous expression in glandular epithelia and neuronal cells. | F-actin cross-linking protein which is thought to anchor actin to a variety of intracellular structures. This is a bundling protein. |
| KIF21A | mid | mid | high | 22 | Cytoplasmic expression in a subset of cells including neuronal cells, respiratory epithelia and cells in seminiferous ducts | Microtubule-binding motor protein probably involved in neuronal axonal transport. In vitro, has a plus-end directed motor activity. |
| MYH9 | low | low | low | 448 | Cytoplasmic and membranous expression in several cell types, mainly in endothelial and glandular cells. | Cellular myosin that appears to play a role in cytokinesis, cell shape, and specialized functions such as secretion and capping. Required for cortical actin clearance prior to oocyte exocytosis |
| SPTBN1 | high | high | mid | 250 | General cytoplasmic and membranous expression. | Interacts with calmodulin and actin, crucial for CNS development and function. |
| DBN1 | high | high | low | 204 | Cytoplasmic and membranous expression in several tissues, most abundant in CNS. | Orchestrates actin polymerization, cell projections, and synaptic plasticity. |
| CAMK2D | mid | high | mid | 52 | Cytoplasmic expression at variable levels in selected tissues, most abundant in heart muscle | Calcium/calmodulin-dependent protein kinase involved in the regulation of Ca(2+) homeostatis and excitation-contraction coupling (ECC) in heart |
| VDAC2 | mid | high | low | 256 | Ubiquitous cytoplasmic expression with a granular pattern. | Forms a channel through the mitochondrial outer membrane that allows diffusion of small hydrophilic molecules |
| CAP2 | low | high | low | 28 | Cytoplasmic expression in the CNS, myocytes and smooth muscle cells. | Involved in the regulation of actin polymerization. |
| ABLIM1 | high | mid | low | 90 | General cytoplasmic and membranous expression. | Encodes a LIM protein mediating interactions between actin and cytoplasmic targets. |
| CALM3*** | high | mid | low | 327 | Ubiquitous cytoplasmic expression | binds calcium and functions as a enzymatic cofactor. Activity of this protein is important in the regulation of the cell cycle and cytokinesis. |
| TPM3 | low | mid | low | 354 | Cytoplasmic expression, mainly in skeletal myocytes. | Binds to actin filaments in muscle and non-muscle cells. |
| LIMCH1 | mid | low | high | 89 | Expression in several tissues. | Activates non-muscle myosin IIa, regulates actin stress fibers, cell spreading, and migration. |
| MYH14 | mid | low | low | 318 | General cytoplasmic and membranous expression, most abundant in gastrointestinal mucosa. | ellular myosin that appears to play a role in cytokinesis, cell shape, and specialized functions such as secretion and capping |
| ANXA1 | low | low | N/A | 68 | Cytoplasmic, membranous and nuclear expression at variable levels in selected tissues. | Regulates immune response, inflammation, and T-cell differentiation; binds calcium and phospholipids. |
| LLGL1 | high | mid | high | 19 | Cytoplasmic expression in most tissues. | Cortical cytoskeleton protein found in a complex involved in maintaining cell polarity and epithelial integrity. I |
| ADD3 | high | high | mid | 120 | Distinct membranous expression in most cell types. | Promotes spectrin-actin network assembly, regulates vascular and renal functions, and supports podocyte structure and function. |
| SPTBN2 | high | high | low | 98 | Cytoplasmic and membranous expression in most cell types, highly expressed in CNS, renal distal tubules and squamous epithelia. | Probably plays an important role in neuronal membrane skeleton. |

| | | | | | | |
|---|---|---|---|---|---|---|
| CTNNB1 | mid | high | mid | 78 | Membranous expression in most tissues. | part of a complex of proteins that constitute adherens junctions . Key downstream component of the canonical Wnt signaling pathway. |
| VDAC1 | low | high | low | 213 | Ubiquitous cytoplasmic expression. | Non-selective ion channel in mitochondrial and plasma membranes, regulates ion transport, apoptosis, mitophagy, and cytochrome c release. |
| MACF1 | high | mid | mid | 113 | Granular cytoplasmic expression | links actin and microtubules |
| IMMT | mid | mid | mid | 93 | General cytoplasmic expression, most abundant in heart, stomach and the renal tubules. | Key MICOS complex component, crucial for maintaining mitochondrial cristae structure and inner membrane architecture. |
| MGST3 | mid | mid | low | 9 | Cytoplasmic expression in several tissues, including muscles, epididymis and gi. | Has glutathione S-transferase and peroxidase activities, detoxifying and synthesizing metabolites by conjugating glutathione to eicosanoids. |
| EFHD2 | mid | mid | low | 139 | Cytoplasmic and membranous expression in several tissues, highest expression in immune cells. | Regulates B-cell apoptosis and negatively controls NF-kappa-B activation. |
| CAPZA2 | mid | mid | low | 232 | Cytoplasmic expression in several tissues | F-actin-capping proteins bind in a Ca(2+)-independent manner to the fast growing ends of actin filaments |
| PLEC | mid | mid | low | 233 | embranous and cytoplasmic expression in almost all cells. | Interlinks intermediate filaments with microtubules and microfilaments and anchors intermediate filaments to desmosomes or hemidesmosomes. |
| TPM4 | mid | mid | low | 311 | General cytoplasmic expression. | Binds actin, regulates striated and smooth muscle contraction, stabilizes cytoskeleton. |
| SYNPO | low | mid | low | 25 | Cytoplasmic expression in several tissues, including skeletal and heart muscle. | Modulates actin dynamics in dendritic spines and podocyte foot processes |
| HSPB1 | low | mid | low | 128 | Cytoplasmic expression mainly in squamous epithelial cells | Small heat shock protein which functions as a molecular chaperone probably maintaining denatured proteins in a folding-competent state |
| MYO1C | low | mid | low | 152 | Cytoplasmic and membranous expression in most tissues. | Actin-based motor protein, involved in glucose transporter recycling and hair cell adaptation. |
| VDAC3 | low | mid | low | 226 | Granular cytoplasmic expression in most tissues. | Forms a mitochondrial outer membrane channel for small molecule diffusion, involved in male fertility and sperm mitochondrial sheath formation. |
| TPM1 | low | low | low | 283 | Cytoplasmic expression mainly in skeletal, heart and smooth muscle cells. | Regulates muscle contraction and stabilizes actin filaments in various cells. |
| EPB41L1 | high | high | high | 83 | Cytoplasmic and membranous expression in most tissues. | May function to confer stability and plasticity to neuronal membrane via multiple interactions, including the spectrin-actin-based cytoskeleton, integral membrane channels and membrane-associated guanylate kinases. |
| PIN1 | high | high | low | 57 | Nuclear and cytoplasmic expression in several tissues with highest levels in neuronal cells of CNS | Peptidyl-prolyl cis/trans isomerases (PPIases) catalyze the cis/trans isomerization of peptidyl-prolyl peptide bonds. This gene encodes one of the PPIases, which specifically binds to phosphorylated ser/thr-pro motifs to catalytically regulate the post-phosphorylation conformation of its substrates. |
| MYO6 | mid | high | high | 148 | Cytoplasmic and membranous expression in most tissues. | Myosins are actin-based motor molecules with ATPase activity |

| | | | | | | |
|---|---|---|---|---|---|---|
| HK1 | mid | high | low | 103 | Ubiquitous cytoplasmic expression with a granular pattern. | Catalyzes the phosphorylation of various hexoses, such as D-glucose, D-glucosamine, D-fructose, D-mannose |
| PHB1 | mid | high | low | 269 | General cytoplasmic expression with a granular pattern. | Protein with pleiotropic attributes mediated in a cell-compartment- and tissue-specific manner, which include the plasma membrane-associated cell signaling functions, mitochondrial chaperone, and transcriptional co-regulator of transcription factors in the nucleus |
| MYO18A | low | high | high | 25 | Ubiquitous cytoplasmic and membranous expression | myosin motor protein |
| CHCHD3 | low | high | high | 61 | General cytoplasmic expression with a granular pattern | Component of the MICOS complex, a large protein complex of the mitochondrial inner membrane that plays crucial roles in the maintenance of crista junctions, inner membrane architecture, and formation of contact sites to the outer membrane |
| SH3BGRL2 | low | high | mid | 19 | Nuclear expression in most tissues. | Predicted to enable SH3 domain binding activity. Located in nuclear membrane and nucleoplasm. |
| SIRT2 | high | mid | high | 5 | Cytoplasmic expression in several tissues, most abundant in skeletal muscle. | protein deacetylase modifies lysines on histones and other proteins, influencing cell cycle, genomic stability, and various biological processes. |
| ADD1 | mid | mid | mid | 140 | Cytoplasmic and membranous expression in most tissues. | Membrane-cytoskeleton-associated protein that promotes the assembly of the spectrin-actin network. Binds to calmodulin. |
| UBXN6 | mid | mid | low | 15 | Cytoplasmic and nuclear expression in most tissues. | The protein may regulate VCP ATPase activity, aiding in lysosomal transport, ERAD, and macroautophagy. |
| ERP44 | mid | mid | low | 64 | Ubiquitous cytoplasmic expression, most abundant in glandular cells. | This ER protein, part of the PDI family, acts as a pH-regulated chaperone, aiding protein quality control. |
| PHB2 | mid | mid | low | 243 | Ubiquitous cytoplasmic expression with a granular pattern. | Enables several functions, including protein C-terminus binding activity; protein N-terminus binding activity; and protein dimerization activity. |
| SORBS1 | low | mid | mid | 17 | General cytoplasmic expression. | Links CBL to insulin receptor, essential for glucose transport and actin stress fiber formation. |
| ACTN2 | low | mid | low | 213 | Cytoplasmic expression in striated muscle and brain. | F-actin cross-linking protein which is thought to anchor actin to a variety of intracellular structures. |
| DST | high | low | high | 84 | Cytoplasmic expression in several tissues, including CNS. | Cytoskeletal linker protein. |
| MYL6 | mid | low | low | 411 | Cytoplasmic expression in several tissues, most abundant in smooth muscle. | Regulatory light chain of myosin. Does not bind calcium. |
| ACTB | mid | low | low | 667 | High myoepithelial expression in all tissues. | Actin beta. Cyotskeleton |
| TMOD1 | low | low | mid | 31 | Expression in smooth and heart muscle and in erythrocytes. | Blocks actin filament elongation and depolymerization at the pointed end. |
| ACTBL2 | N/A | N/A | N/A | 628 | Cytoplasmic expression in lymphoid cells and smooth muscle. | Actins are highly conserved proteins that are involved in various types of cell motility and are ubiquitously expressed in all eukaryotic cells. |

** OGs = oligodendrocytes ** CRAPome score: 0-716, indicating how often the protein was detected as a nonspecific hit in 716 independent IP experiments using standardized negative controls on human tissues *** CALM3;CALM2;CALM1*
